## Supplementary material for "Building Transformers from Neurons and Astrocytes": LaTeX files: bio_transformer.bib

@article{amari1998natural,
  title={Natural gradient works efficiently in learning},
  author={Amari, Shun-Ichi},
  journal={Neural computation},
  volume={10},
  number={2},
  pages={251--276},
  year={1998},
  publisher={MIT Press}
}


@inproceedings{sander2022sinkformers,
  title={Sinkformers: Transformers with doubly stochastic attention},
  author={Sander, Michael E and Ablin, Pierre and Blondel, Mathieu and Peyr{\'e}, Gabriel},
  booktitle={International Conference on Artificial Intelligence and Statistics},
  pages={3515--3530},
  year={2022},
  organization={PMLR}
}

@inproceedings{katharopoulos2020transformers,
  title={Transformers are rnns: Fast autoregressive transformers with linear attention},
  author={Katharopoulos, Angelos and Vyas, Apoorv and Pappas, Nikolaos and Fleuret, Fran{\c{c}}ois},
  booktitle={International Conference on Machine Learning},
  pages={5156--5165},
  year={2020},
  organization={PMLR}
}

@article{toneva2019interpreting,
  title={Interpreting and improving natural-language processing (in machines) with natural language-processing (in the brain)},
  author={Toneva, Mariya and Wehbe, Leila},
  journal={Advances in Neural Information Processing Systems},
  volume={32},
  year={2019}
}

@article{caucheteux2022brains,
  title={Brains and algorithms partially converge in natural language processing},
  author={Caucheteux, Charlotte and King, Jean-R{\'e}mi},
  journal={Communications biology},
  volume={5},
  number={1},
  pages={1--10},
  year={2022},
  publisher={Nature Publishing Group}
}

@article{bricken2021attention,
  title={Attention approximates sparse distributed memory},
  author={Bricken, Trenton and Pehlevan, Cengiz},
  journal={Advances in Neural Information Processing Systems},
  volume={34},
  pages={15301--15315},
  year={2021}
}


 @article{Whittington_Warren_Behrens_2022, title={Relating transformers to models and neural representations of the hippocampal formation}, url={http://arxiv.org/abs/2112.04035}, abstractNote={Many deep neural network architectures loosely based on brain networks have recently been shown to replicate neural firing patterns observed in the brain. One of the most exciting and promising novel architectures, the Transformer neural network, was developed without the brain in mind. In this work, we show that transformers, when equipped with recurrent position encodings, replicate the precisely tuned spatial representations of the hippocampal formation; most notably place and grid cells. Furthermore, we show that this result is no surprise since it is closely related to current hippocampal models from neuroscience. We additionally show the transformer version offers dramatic performance gains over the neuroscience version. This work continues to bind computations of artificial and brain networks, offers a novel understanding of the hippocampal-cortical interaction, and suggests how wider cortical areas may perform complex tasks beyond current neuroscience models such as language comprehension.}, note={arXiv: 2112.04035}, journal={arXiv:2112.04035 [cs, q-bio]}, author={Whittington, James C. R. and Warren, Joseph and Behrens, Timothy E. J.}, year={2022}, month={Mar} }


@article{schrimpf2021neural,
  title={The neural architecture of language: Integrative modeling converges on predictive processing},
  author={Schrimpf, Martin and Blank, Idan Asher and Tuckute, Greta and Kauf, Carina and Hosseini, Eghbal A and Kanwisher, Nancy and Tenenbaum, Joshua B and Fedorenko, Evelina},
  journal={Proceedings of the National Academy of Sciences},
  volume={118},
  number={45},
  pages={e2105646118},
  year={2021},
  publisher={National Acad Sciences}
}

 @inproceedings{Vaswani_Shazeer_Parmar_Uszkoreit_Jones_Gomez_Kaiser_Polosukhin_2017, title={Attention is All you Need}, volume={30}, url={https://proceedings.neurips.cc/paper/2017/hash/3f5ee243547dee91fbd053c1c4a845aa-Abstract.html}, booktitle={Advances in Neural Information Processing Systems}, publisher={Curran Associates, Inc.}, author={Vaswani, Ashish and Shazeer, Noam and Parmar, Niki and Uszkoreit, Jakob and Jones, Llion and Gomez, Aidan N and Kaiser, Łukasz and Polosukhin, Illia}, year={2017} }


@article{croft2015plasticity,
  title={Plasticity of neuron-glial transmission: equipping glia for long-term integration of network activity},
  author={Croft, Wayne and Dobson, Katharine L and Bellamy, Tomas C},
  journal={Neural plasticity},
  volume={2015},
  year={2015},
  publisher={Hindawi}
}

@article{stobart2018cortical,
  title={Cortical circuit activity evokes rapid astrocyte calcium signals on a similar timescale to neurons},
  author={Stobart, Jillian L and Ferrari, Kim David and Barrett, Matthew JP and Gl{\"u}ck, Chaim and Stobart, Michael J and Zuend, Marc and Weber, Bruno},
  journal={Neuron},
  volume={98},
  number={4},
  pages={726--735},
  year={2018},
  publisher={Elsevier}
}

 @article{Lind_Brazhe_Jessen_Tan_Lauritzen_2013, title={Rapid stimulus-evoked astrocyte Ca2+ elevations and hemodynamic responses in mouse somatosensory cortex in vivo}, volume={110}, DOI={10.1073/pnas.1310065110}, number={48}, journal={Proceedings of the National Academy of Sciences}, publisher={Proceedings of the National Academy of Sciences}, author={Lind, Barbara Lykke and Brazhe, Alexey R. and Jessen, Sanne Barsballe and Tan, Florence C. C. and Lauritzen, Martin J.}, year={2013}, month={Nov}, pages={E4678–E4687} }


@article{klampfl2013emergence,
  title={Emergence of dynamic memory traces in cortical microcircuit models through STDP},
  author={Klampfl, Stefan and Maass, Wolfgang},
  journal={Journal of Neuroscience},
  volume={33},
  number={28},
  pages={11515--11529},
  year={2013},
  publisher={Soc Neuroscience}
}

 @article{Yamins_Hong_Cadieu_Solomon_Seibert_DiCarlo_2014, title={Performance-optimized hierarchical models predict neural responses in higher visual cortex}, volume={111}, DOI={10.1073/pnas.1403112111}, number={23}, journal={Proceedings of the National Academy of Sciences}, publisher={Proceedings of the National Academy of Sciences}, author={Yamins, Daniel L. K. and Hong, Ha and Cadieu, Charles F. and Solomon, Ethan A. and Seibert, Darren and DiCarlo, James J.}, year={2014}, month={Jun}, pages={8619–8624} }


 @inproceedings{Katharopoulos_Vyas_Pappas_Fleuret_2020, title={Transformers are RNNs: Fast Autoregressive Transformers with Linear Attention}, ISSN={2640-3498}, url={https://proceedings.mlr.press/v119/katharopoulos20a.html}, abstractNote={Transformers achieve remarkable performance in several tasks but due to their quadratic complexity, with respect to the input’s length, they are prohibitively slow for very long sequences. To address this limitation, we express the self-attention as a linear dot-product of kernel feature maps and make use of the associativity property of matrix products to reduce the complexity from $bigO{N^2}$ to $bigO{N}$, where $N$ is the sequence length. We show that this formulation permits an iterative implementation that dramatically accelerates autoregressive transformers and reveals their relationship to recurrent neural networks. Our emph{Linear Transformers} achieve similar performance to vanilla Transformers and they are up to 4000x faster on autoregressive prediction of very long sequences.}, booktitle={Proceedings of the 37th International Conference on Machine Learning}, publisher={PMLR}, author={Katharopoulos, Angelos and Vyas, Apoorv and Pappas, Nikolaos and Fleuret, François}, year={2020}, month={Nov}, pages={5156–5165}, language={en} }


 @article{Lu_Li_He_Sun_Dong_Qin_Wang_Liu_2019, title={Understanding and Improving Transformer From a Multi-Particle Dynamic System Point of View}, url={http://arxiv.org/abs/1906.02762}, abstractNote={The Transformer architecture is widely used in natural language processing. Despite its success, the design principle of the Transformer remains elusive. In this paper, we provide a novel perspective towards understanding the architecture: we show that the Transformer can be mathematically interpreted as a numerical Ordinary Differential Equation (ODE) solver for a convection-diffusion equation in a multi-particle dynamic system. In particular, how words in a sentence are abstracted into contexts by passing through the layers of the Transformer can be interpreted as approximating multiple particles’ movement in the space using the Lie-Trotter splitting scheme and the Euler’s method. Given this ODE’s perspective, the rich literature of numerical analysis can be brought to guide us in designing effective structures beyond the Transformer. As an example, we propose to replace the Lie-Trotter splitting scheme by the Strang-Marchuk splitting scheme, a scheme that is more commonly used and with much lower local truncation errors. The Strang-Marchuk splitting scheme suggests that the self-attention and position-wise feed-forward network (FFN) sub-layers should not be treated equally. Instead, in each layer, two position-wise FFN sub-layers should be used, and the self-attention sub-layer is placed in between. This leads to a brand new architecture. Such an FFN-attention-FFN layer is “Macaron-like”, and thus we call the network with this new architecture the Macaron Net. Through extensive experiments, we show that the Macaron Net is superior to the Transformer on both supervised and unsupervised learning tasks. The reproducible codes and pretrained models can be found at https://github.com/zhuohan123/macaron-net}, note={arXiv: 1906.02762}, journal={arXiv:1906.02762 [cs, stat]}, author={Lu, Yiping and Li, Zhuohan and He, Di and Sun, Zhiqing and Dong, Bin and Qin, Tao and Wang, Liwei and Liu, Tie-Yan}, year={2019}, month={Jun}, language={en} }


 @article{Peterson_2022, title={What can astrocytes compute?}, url={https://www.biorxiv.org/content/early/2022/01/12/2021.10.20.465192}, DOI={10.1101/2021.10.20.465192}, abstractNote={A foundational result in neural computation was proving the firing-rate model of neurons defines a universal function approximator. In this paper I prove it is possible for astrocytes to compute anything neurons can, by extending this original proof to a model of calcium waves in astrocytes. I confirm this in a series of computer simulations. The major limit for astrocytes, it turns out, is not their ability to learn solutions but the computational complexity of their network. I suggest some initial experiments that might be used to confirm these predictions.Competing Interest StatementThe authors have declared no competing interest.}, journal={bioRxiv}, publisher={Cold Spring Harbor Laboratory}, author={Peterson, Erik J.}, year={2022} }


@inproceedings{tang2019introducing,
  title={Introducing astrocytes on a neuromorphic processor: Synchronization, local plasticity and edge of chaos},
  author={Tang, Guangzhi and Polykretis, Ioannis E and Ivanov, Vladimir A and Shah, Arpit and Michmizos, Konstantinos P},
  booktitle={Proceedings of the 7th Annual Neuro-inspired Computational Elements Workshop},
  pages={1--9},
  year={2019}
}

@article{ivanov2021increasing,
  title={Increasing Liquid State Machine Performance with Edge-of-Chaos Dynamics Organized by Astrocyte-modulated Plasticity},
  author={Ivanov, Vladimir and Michmizos, Konstantinos},
  journal={Advances in Neural Information Processing Systems},
  volume={34},
  pages={25703--25719},
  year={2021}
}

 @article{Perea_Navarrete_Araque_2009, title={Tripartite synapses: astrocytes process and control synaptic information}, volume={32}, ISSN={0166-2236}, DOI={10.1016/j.tins.2009.05.001}, abstractNote={The term ‘tripartite synapse’ refers to a concept in synaptic physiology based on the demonstration of the existence of bidirectional communication between astrocytes and neurons. Consistent with this concept, in addition to the classic ‘bipartite’ information flow between the pre- and postsynaptic neurons, astrocytes exchange information with the synaptic neuronal elements, responding to synaptic activity and, in turn, regulating synaptic transmission. Because recent evidence has demonstrated that astrocytes integrate and process synaptic information and control synaptic transmission and plasticity, astrocytes, being active partners in synaptic function, are cellular elements involved in the processing, transfer and storage of information by the nervous system. Consequently, in contrast to the classically accepted paradigm that brain function results exclusively from neuronal activity, there is an emerging view, which we review herein, in which brain function actually arises from the coordinated activity of a network comprising both neurons and glia.}, number={8}, journal={Trends in Neurosciences}, author={Perea, Gertrudis and Navarrete, Marta and Araque, Alfonso}, year={2009}, month={Aug}, pages={421–431}, language={en} }


@article{rahimi2007random,
  title={Random features for large-scale kernel machines},
  author={Rahimi, Ali and Recht, Benjamin},
  journal={Advances in neural information processing systems},
  volume={20},
  year={2007}
}

@book{verkhratsky2007glial,
  title={Glial neurobiology: a textbook},
  author={Verkhratsky, Alexei and Butt, Arthur},
  year={2007},
  publisher={John Wiley \& Sons}
}

@article{scemes2006astrocyte,
  title={Astrocyte calcium waves: what they are and what they do},
  author={Scemes, Eliana and Giaume, Christian},
  journal={Glia},
  volume={54},
  number={7},
  pages={716--725},
  year={2006},
  publisher={Wiley Online Library}
}

@article{peng2021random,
  title={Random feature attention},
  author={Peng, Hao and Pappas, Nikolaos and Yogatama, Dani and Schwartz, Roy and Smith, Noah A and Kong, Lingpeng},
  journal={arXiv preprint arXiv:2103.02143},
  year={2021}
}

 @article{Kuga_Sasaki_Takahara_Matsuki_Ikegaya_2011, title={Large-Scale Calcium Waves Traveling through Astrocytic Networks In Vivo}, volume={31}, ISSN={0270-6474}, DOI={10.1523/JNEUROSCI.5319-10.2011}, abstractNote={Macroscopic changes in cerebral blood flow, such as those captured by functional imaging of the brain, require highly organized, large-scale dynamics of astrocytes, glial cells that interact with both neuronal and cerebrovascular networks. However, astrocyte activity has been studied mainly at the level of individual cells, and information regarding their collective behavior is lacking. In this work, we monitored calcium activity simultaneously from hundreds of mouse hippocampal astrocytes in vivo and found that almost all astrocytes participated en masse in regenerative waves that propagated from cell to cell (referred to here as “glissandi”). Glissandi emerged depending on the neuronal activity and accompanied a reduction in infraslow fluctuations of local field potentials and a decrease in the flow of red blood cells. This novel phenomenon was heretofore overlooked, probably because of the high vulnerability of astrocytes to light damage; glissandi occurred only when observed at much lower laser intensities than previously used.}, number={7}, journal={The Journal of Neuroscience}, author={Kuga, Nahoko and Sasaki, Takuya and Takahara, Yuji and Matsuki, Norio and Ikegaya, Yuji}, year={2011}, month={Feb}, pages={2607–2614} }
 
 
@article{lin2021survey,
  title={A survey of transformers},
  author={Lin, Tianyang and Wang, Yuxin and Liu, Xiangyang and Qiu, Xipeng},
  journal={arXiv preprint arXiv:2106.04554},
  year={2021}
}
 
@book{kandel2000principles,
  title={Principles of neural science},
  author={Kandel, Eric R and Schwartz, James H and Jessell, Thomas M and Siegelbaum, Steven and Hudspeth, A James and Mack, Sarah and others},
  volume={4},
  year={2000},
  publisher={McGraw-hill New York}
}

 @article{Sul_Orosz_Givens_Haydon_2004, title={Astrocytic Connectivity in the Hippocampus}, volume={1}, ISSN={1740-925X}, DOI={10.1017/s1740925x04000031}, abstractNote={Little is known about the functional connectivity between astrocytes in the CNS. To explore this issue we photo-released glutamate onto a single astrocyte in murine hippocampal slices and imaged calcium responses. Photo-release of glutamate causes a metabotropic glutamate receptor (mGluR)-dependent increase in internal calcium in the stimulated astrocyte and delayed calcium elevations in neighboring cells. The delayed elevation in calcium was not caused by either neuronal activity following synaptic transmission or by glutamate released from astrocytes. However, it was reduced by flufenamic acid (FFA), which is consistent with a role for adenosine triphosphate (ATP) release from astrocytes as an intercellular messenger. Exogenous ligands such as ATP (1 μM) increased the number of astrocytes that were recruited into coupled astrocytic networks, indicating that extracellular accumulation of neurotransmitters modulates neuronal excitability, synaptic transmission and functional coupling between astrocytes.}, number={1}, journal={Neuron glia biology}, author={Sul, Jai-Yoon and Orosz, George and Givens, Richard S. and Haydon, Philip G.}, year={2004}, month={Feb}, pages={3–11} }
 
 
  @article{Newman_2003, title={New roles for astrocytes: Regulation of synaptic transmission}, volume={26}, ISSN={0166-2236}, DOI={10.1016/S0166-2236(03)00237-6}, abstractNote={Although glia often envelop synapses, they have traditionally been viewed as passive participants in synaptic function. Recent evidence has demonstrated, however, that there is a dynamic two-way communication between glia and neurons at the synapse. Neurotransmitters released from presynaptic neurons evoke Ca2+ concentration increases in adjacent glia. Activated glia, in turn, release transmitters, including glutamate and ATP. These gliotransmitters feed back onto the presynaptic terminal either to enhance or to depress further release of neurotransmitter. Transmitters released from glia can also directly stimulate postsynaptic neurons, producing either excitatory or inhibitory responses. Based on these new findings, glia should be considered an active partner at the synapse, dynamically regulating synaptic transmission.}, number={10}, journal={Trends in Neurosciences}, author={Newman, Eric A}, year={2003}, month={Oct}, pages={536–542}, language={en} }
  
  @article{Gordon_Iremonger_Kantevari_Ellis-Davies_MacVicar_Bains_2009, title={Astrocyte-Mediated Distributed Plasticity at Hypothalamic Glutamate Synapses}, volume={64}, ISSN={08966273}, DOI={10.1016/j.neuron.2009.10.021}, abstractNote={Afferent activity can induce fast, feed-forward changes in synaptic efﬁcacy that are synapse speciﬁc. Using combined electrophysiology, caged molecule photolysis, and Ca2+ imaging, we describe a plasticity in which the recruitment of astrocytes in response to afferent activity causes a fast and feed-forward, yet distributed increase in the amplitude of quantal synaptic currents at multiple glutamate synapses on magnocellular neurosecretory cells in the hypothalamic paraventricular nucleus. The plasticity is largely multiplicative, consistent with a proportional increase or ‘“scaling”’ in the strength of all synapses on the neuron. This effect requires a metabotropic glutamate receptor-mediated rise in Ca2+ in the astrocyte processes surrounding the neuron and the release of the gliotransmitter ATP, which acts on postsynaptic purinergic receptors. These data provide evidence for a form of distributed synaptic plasticity that is feed-forward, expressed quickly, and mediated by the synaptic activation of neighboring astrocytes.}, number={3}, journal={Neuron}, author={Gordon, Grant R.J. and Iremonger, Karl J. and Kantevari, Srinivas and Ellis-Davies, Graham C.R. and MacVicar, Brian A. and Bains, Jaideep S.}, year={2009}, month={Nov}, pages={391–403}, language={en} }


 @article{yang2022transformers,
  title={Transformers from an Optimization Perspective},
  author={Yang, Yongyi and Huang, Zengfeng and Wipf, David},
  journal={arXiv preprint arXiv:2205.13891},
  year={2022}
}

@article{rasmusson2000role,
  title={The role of acetylcholine in cortical synaptic plasticity},
  author={Rasmusson, DD},
  journal={Behavioural brain research},
  volume={115},
  number={2},
  pages={205--218},
  year={2000},
  publisher={Elsevier}
}

@article{escartin2021reactive,
  title={Reactive astrocyte nomenclature, definitions, and future directions},
  author={Escartin, Carole and Galea, Elena and Lakatos, Andr{\'a}s and O’Callaghan, James P and Petzold, Gabor C and Serrano-Pozo, Alberto and Steinh{\"a}user, Christian and Volterra, Andrea and Carmignoto, Giorgio and Agarwal, Amit and others},
  journal={Nature neuroscience},
  volume={24},
  number={3},
  pages={312--325},
  year={2021},
  publisher={Nature Publishing Group}
}

@article{semyanov2021astrocytic,
  title={Astrocytic processes: from tripartite synapses to the active milieu},
  author={Semyanov, Alexey and Verkhratsky, Alexei},
  journal={Trends in neurosciences},
  volume={44},
  number={10},
  pages={781--792},
  year={2021},
  publisher={Elsevier}
}

@article{schrimpf2020brain,
  title={Brain-score: Which artificial neural network for object recognition is most brain-like?},
  author={Schrimpf, Martin and Kubilius, Jonas and Hong, Ha and Majaj, Najib J and Rajalingham, Rishi and Issa, Elias B and Kar, Kohitij and Bashivan, Pouya and Prescott-Roy, Jonathan and Geiger, Franziska and others},
  journal={BioRxiv},
  pages={407007},
  year={2020},
  publisher={Cold Spring Harbor Laboratory}
}

@article{hopfield1982neural,
  title={Neural networks and physical systems with emergent collective computational abilities.},
  author={Hopfield, John J},
  journal={Proceedings of the national academy of sciences},
  volume={79},
  number={8},
  pages={2554--2558},
  year={1982},
  publisher={National Acad Sciences}
}


 @inproceedings{Tyulmankov_Fang_Vadaparty_Yang_2021, title={Biological learning in key-value memory networks}, volume={34}, url={https://proceedings.neurips.cc/paper/2021/hash/bacadc62d6e67d7897cef027fa2d416c-Abstract.html}, abstractNote={In neuroscience, classical Hopfield networks are the standard biologically plausible model of long-term memory, relying on Hebbian plasticity for storage and attractor dynamics for recall. In contrast, memory-augmented neural networks in machine learning commonly use a key-value mechanism to store and read out memories in a single step. Such augmented networks achieve impressive feats of memory compared to traditional variants, yet their biological relevance is unclear. We propose an implementation of basic key-value memory that stores inputs using a combination of biologically plausible three-factor plasticity rules. The same rules are recovered when network parameters are meta-learned. Our network performs on par with classical Hopfield networks on autoassociative memory tasks and can be naturally extended to continual recall, heteroassociative memory, and sequence learning. Our results suggest a compelling alternative to the classical Hopfield network as a model of biological long-term memory.}, booktitle={Advances in Neural Information Processing Systems}, publisher={Curran Associates, Inc.}, author={Tyulmankov, Danil and Fang, Ching and Vadaparty, Annapurna and Yang, Guangyu Robert}, year={2021}, pages={22247–22258} }
 
 
 
@article{butts2006tuning,
  title={Tuning curves, neuronal variability, and sensory coding},
  author={Butts, Daniel A and Goldman, Mark S},
  journal={PLoS biology},
  volume={4},
  number={4},
  pages={e92},
  year={2006},
  publisher={Public Library of Science San Francisco, USA}
}


 @article{Oberheim_2009, title={Uniquely Hominid Features of Adult Human Astrocytes}, volume={29}, ISSN={0270-6474, 1529-2401}, DOI={10.1523/JNEUROSCI.4707-08.2009}, number={10}, journal={Journal of Neuroscience}, author={Oberheim, N. A. and Takano, T. and Han, X. and He, W. and Lin, J. H. C. and Wang, F. and Xu, Q. and Wyatt, J. D. and Pilcher, W. and Ojemann, J. G. and Ransom, B. R. and Goldman, S. A. and Nedergaard, M.}, year={2009}, month={Mar}, pages={3276–3287}, language={en} }

@article{halassa2007synaptic,
  title={Synaptic islands defined by the territory of a single astrocyte},
  author={Halassa, Michael M and Fellin, Tommaso and Takano, Hajime and Dong, Jing-Hui and Haydon, Philip G},
  journal={Journal of Neuroscience},
  volume={27},
  number={24},
  pages={6473--6477},
  year={2007},
  publisher={Soc Neuroscience}
}

@article{von2016search,
  title={The search for true numbers of neurons and glial cells in the human brain: A review of 150 years of cell counting},
  author={Von Bartheld, Christopher S and Bahney, Jami and Herculano-Houzel, Suzana},
  journal={Journal of Comparative Neurology},
  volume={524},
  number={18},
  pages={3865--3895},
  year={2016},
  publisher={Wiley Online Library}
}

@article{song2016design,
  title={Design of logic gates using spiking neural P systems with homogeneous neurons and astrocytes-like control},
  author={Song, Tao and Zheng, Pan and Wong, ML Dennis and Wang, Xun},
  journal={Information Sciences},
  volume={372},
  pages={380--391},
  year={2016},
  publisher={Elsevier}
}

@article{kastanenka2020roadmap,
  title={A roadmap to integrate astrocytes into Systems Neuroscience},
  author={Kastanenka, Ksenia V and Moreno-Bote, Rub{\'e}n and De Pitt{\`a}, Maurizio and Perea, Gertrudis and Eraso-Pichot, Abel and Masgrau, Roser and Poskanzer, Kira E and Galea, Elena},
  journal={Glia},
  volume={68},
  number={1},
  pages={5--26},
  year={2020},
  publisher={Wiley Online Library}
}

@article{de2012computational,
  title={Computational quest for understanding the role of astrocyte signaling in synaptic transmission and plasticity},
  author={De Pitt{\`a}, Maurizio and Volman, Vladislav and Berry, Hugues and Parpura, Vladimir and Volterra, Andrea and Ben-Jacob, Eshel},
  journal={Frontiers in computational neuroscience},
  volume={6},
  pages={98},
  year={2012},
  publisher={Frontiers Media SA}
}

@inproceedings{krotov2020large,
  title={Large Associative Memory Problem in Neurobiology and Machine Learning},
  author={Krotov, Dmitry and Hopfield, John J},
  booktitle={International Conference on Learning Representations},
  year={2020}
}


@article{carandini2012normalization,
  title={Normalization as a canonical neural computation},
  author={Carandini, Matteo and Heeger, David J},
  journal={Nature Reviews Neuroscience},
  volume={13},
  number={1},
  pages={51--62},
  year={2012},
  publisher={Nature Publishing Group}
}

@article{de2009glutamate,
  title={Glutamate regulation of calcium and IP3 oscillating and pulsating dynamics in astrocytes},
  author={De Pitt{\`a}, Maurizio and Goldberg, Mati and Volman, Vladislav and Berry, Hugues and Ben-Jacob, Eshel},
  journal={Journal of biological physics},
  volume={35},
  number={4},
  pages={383--411},
  year={2009},
  publisher={Springer}
}

@article{de2011tale,
  title={A tale of two stories: astrocyte regulation of synaptic depression and facilitation},
  author={De Pitt{\`a}, Maurizio and Volman, Vladislav and Berry, Hugues and Ben-Jacob, Eshel},
  journal={PLoS computational biology},
  volume={7},
  number={12},
  pages={e1002293},
  year={2011},
  publisher={Public Library of Science San Francisco, USA}
}

@article{goldberg2010nonlinear,
  title={Nonlinear gap junctions enable long-distance propagation of pulsating calcium waves in astrocyte networks},
  author={Goldberg, Mati and De Pitt{\`a}, Maurizio and Volman, Vladislav and Berry, Hugues and Ben-Jacob, Eshel},
  journal={PLoS computational biology},
  volume={6},
  number={8},
  pages={e1000909},
  year={2010},
  publisher={Public Library of Science San Francisco, USA}
}

@article{chance2000divisive,
  title={Divisive inhibition in recurrent networks},
  author={Chance, Frances S and Abbott, LF},
  journal={Network: Computation in Neural Systems},
  volume={11},
  number={2},
  pages={119},
  year={2000},
  publisher={IOP Publishing}
}

 @article{Shen_Wang_Navlakha_2021, title={A Correspondence Between Normalization Strategies in Artificial and Biological Neural Networks}, volume={33}, ISSN={0899-7667, 1530-888X}, DOI={10.1162/neco_a_01439}, abstractNote={Abstract
            A fundamental challenge at the interface of machine learning and neuroscience is to uncover computational principles that are shared between artificial and biological neural networks. In deep learning, normalization methods such as batch normalization, weight normalization, and their many variants help to stabilize hidden unit activity and accelerate network training, and these methods have been called one of the most important recent innovations for optimizing deep networks. In the brain, homeostatic plasticity represents a set of mechanisms that also stabilize and normalize network activity to lie within certain ranges, and these mechanisms are critical for maintaining normal brain function. In this article, we discuss parallels between artificial and biological normalization methods at four spatial scales: normalization of a single neuron’s activity, normalization of synaptic weights of a neuron, normalization of a layer of neurons, and normalization of a network of neurons. We argue that both types of methods are functionally equivalent—that is, both push activation patterns of hidden units toward a homeostatic state, where all neurons are equally used—and we argue that such representations can improve coding capacity, discrimination, and regularization. As a proof of concept, we develop an algorithm, inspired by a neural normalization technique called synaptic scaling, and show that this algorithm performs competitively against existing normalization methods on several data sets. Overall, we hope this bidirectional connection will inspire neuroscientists and machine learners in three ways: to uncover new normalization algorithms based on established neurobiological principles; to help quantify the trade-offs of different homeostatic plasticity mechanisms used in the brain; and to offer insights about how stability may not hinder, but may actually promote, plasticity.}, number={12}, journal={Neural Computation}, author={Shen, Yang and Wang, Julia and Navlakha, Saket}, year={2021}, month={Nov}, pages={3179–3203}, language={en} }


 @article{Kofuji_Araque_2021, title={Astrocytes and Behavior}, volume={44}, ISSN={0147-006X, 1545-4126}, DOI={10.1146/annurev-neuro-101920-112225}, abstractNote={Animal behavior was classically considered to be determined exclusively by neuronal activity, whereas surrounding glial cells such as astrocytes played only supportive roles. However, astrocytes are as numerous as neurons in the mammalian brain, and current findings indicate a chemically based dialog between astrocytes and neurons. Activation of astrocytes by synaptically released neurotransmitters converges on regulating intracellular Ca2+ in astrocytes, which then can regulate the efficacy of near and distant tripartite synapses at diverse timescales through gliotransmitter release. Here, we discuss recent evidence on how diverse behaviors are impacted by this dialog. These recent findings support a paradigm shift in neuroscience, in which animal behavior does not result exclusively from neuronal activity but from the coordinated activity of both astrocytes and neurons. Decoding how astrocytes and neurons interact with each other in various brain circuits will be fundamental to fully understanding how behaviors originate and become dysregulated in disease.}, number={1}, journal={Annual Review of Neuroscience}, author={Kofuji, Paulo and Araque, Alfonso}, year={2021}, month={Jul}, pages={49–67}, language={en} }

@article{de2016astrocytes,
  title={Astrocytes: Orchestrating synaptic plasticity?},
  author={De Pitt{\`a}, Maurizio and Brunel, Nicolas and Volterra, Andrea},
  journal={Neuroscience},
  volume={323},
  pages={43--61},
  year={2016},
  publisher={Elsevier}
}

@article{perea2007astrocytes,
  title={Astrocytes potentiate transmitter release at single hippocampal synapses},
  author={Perea, Gertrudis and Araque, Alfonso},
  journal={Science},
  volume={317},
  number={5841},
  pages={1083--1086},
  year={2007},
  publisher={American Association for the Advancement of Science}
}

@article{perea2009tripartite,
  title={Tripartite synapses: astrocytes process and control synaptic information},
  author={Perea, Gertrudis and Navarrete, Marta and Araque, Alfonso},
  journal={Trends in neurosciences},
  volume={32},
  number={8},
  pages={421--431},
  year={2009},
  publisher={Elsevier}
}

@article{lopez2019astrocyte,
  title={Astrocyte subdomains respond independently in vivo},
  author={L{\'o}pez-Hidalgo, M{\'o}nica and Kellner, Vered and Schummers, James},
  journal={bioRxiv},
  pages={675769},
  year={2019},
  publisher={Cold Spring Harbor Laboratory}
}
