## Supplementary material for "Building Transformers from Neurons and Astrocytes": LaTeX files: extras.txt

\section{Introduction}
Transformer blocks consist of this self-attention mechanism combined with a residual connection and an additional MLP. We will ignore the MLP for now.  Given an embedding matrix $X \in \fR^{N \times \edim}$, self-attention is a map from $\fR^{N \times \edim}$ to $\fR^{N \times \edim}$. Here $N$ is the number of elements in the embedded sequence, and $\edim$ is the embedding dimension.

\[A(X) \coloneqq \softmax\Bigg(\frac{X W_Q W^T_K X^T}{\sqrt{\edim}}\Bigg)XW_V\]
where $W_Q, W_K, W_V$ are the query, key, and value matrices. It will be useful later on to write the softmax function as the product of two matrices:

\[\text{softmax}(Y) = G^{-1}(Y)\text{exp}(Y) \]
where $\text{exp}(\cdot)$ denotes elementwise exponentiation, and $G(Y)$ is a diagonal, positive matrix that normalizes the rows of $\text{exp}(Y)$:

\[G(Y)_{ii} = \sum_{j}\text{exp}(Y_{ij})  \]
Tranformer blocks are often chained together into layers.

\[X_{i+1} = X_{i} + A(X_i) \]
Which may be seen as an Euler integration with unit step-size of the following neural ODE:

\begin{equation}\label{eq:trans_ode}
\dt{X}= A(X)    
\end{equation}

\section{Energy Function}
We would like to know when there is some energy function $E(X) : \fR^{N \times D} \rightarrow \fR$ such that $E(X)$ is always non-increasing along trajectories of \eqref{eq:trans_ode}. Consider the matrix 

\[P(X) \coloneqq \text{exp}(XBB^TX^T) \]
where $B$ is an arbitrary matrix. We will show that the function:

\begin{equation}\label{eq:energy_func}
E \coloneqq \sum_{i,j = 1} P_{ij}(X) \geq 0 
\end{equation}
satisfies the above requirement. Differentiating \eqref{eq:energy_func} with respect to $X$ directly yields:

\[\frac{\partial E}{\partial X} = 2\text{exp}(XBB^TX^T)XBB^T \]
If we tie the query, key, and value weights such that:
\begin{equation}\label{eq:weight_tying}
\frac{W_Q W^T_K}{\sqrt{D}} = BB^T \hspace{1cm} W_V = -2BB^T 
\end{equation}
then we can see that the Transformer ODE \eqref{eq:trans_ode} can be written as:
\begin{equation}\label{eq:natural_gradient_flow}
\dt{X} = A(X) = -G^{-1}(X)\frac{\partial E}{\partial X}
\end{equation}
where $G$ is the normalizing matrix for the softmax function, described above. The weight-tying condition \eqref{eq:weight_tying} is similar to the one described in \cite{sander2022sinkformers} (Assumption 1). The structure of equation \eqref{eq:natural_gradient_flow} is reminiscent of natural gradient flows \citep{amari1998natural}. To show that $E$ is non-increasing along  trajectories of \eqref{eq:natural_gradient_flow}, we directly evaluate the time-derivative:

\[\dt{E} = \sum_{i,j = 1} \frac{\partial E}{\partial X_{ij}} \frac{\partial X_{ij}}{\partial t} = -\sum_{i,j = 1} \frac{1}{G(X)_{ii}}\bigg(\frac{\partial E}{\partial X_{ij}}\bigg)^2 \leq 0   \]
which proves the result. A Google Colab notebook to play with this result is available \href{https://colab.research.google.com/drive/1zFtaz8DymFDlukKYIZKMf5FCIarLwc03?usp=sharing}{here}.

%\includegraphics[scale=0.7]{Energy_vs_Time.png}


This follows from a straightforward calculation:

\begin{align}
\begin{split}
\phi^T_{dot}(x)\phi_{dot}(y) = \phi_{rbf}^T(x)\text{exp}(\frac{||x||^2}{2})\phi_{rbf}(y)\text{exp}(\frac{||y||^2}{2}) \\ \approx 
\text{RBF}(x,y)\text{exp}(\frac{||x||^2}{2})\text{exp}(\frac{||y||^2}{2}) \\ = 
\text{exp}(x^Ty)
\end{split}
\end{align}

A tremendous amount has been learned about glial cells in the last few decades. Despite this, the jury is still out on many basic questions of glial biology. It was originally thought that glial cells (in particular astrocytes) only operate on much slow timescales than neurons. However, recent work has shown fast calcium responses in astrocyte processes on the order of just a few hundred milliseconds \citep{stobart2018cortical}. These details are crucial for computational neuroscientists, as they define the space of `permissible' operations within a particular circuit model. We have tried to distill the important facts of glial biology into the simplest conceivable model. 
\lk{Is there evidence for reading and writing operations?}
