## Supplementary material for "Building Transformers from Neurons and Astrocytes": LaTeX files: main.tex

\RequirePackage[2020-02-02]{latexrelease}
\documentclass[times, twoside]{zHenriquesLab-StyleBioRxiv}
\usepackage{blindtext}
\usepackage{amsmath}
\usepackage{authblk}
\usepackage{amsfonts}
\usepackage{mathtools}
\usepackage{accents}
\usepackage{graphicx}
\usepackage{amssymb}
\usepackage{xcolor}

\newcommand{\lk}[1]{\textcolor{red}{Leo: #1}}
\newcommand{\dk}[1]{\textcolor{blue}{Dima: #1}}
\input{preamble}

% Please give the surname of the lead author for the running footer
\leadauthor{Henriques} 

\begin{document}

\title{Building Transformers from \\ Neurons and Astrocytes}
\shorttitle{Building Transformers from Neurons and Glia}

% Use letters for affiliations, numbers to show equal authorship (if applicable) and to indicate the corresponding author
\author[a,b,\footnote{Work completed during MIT-IBM Watson AI Lab Summer 2022 Internship.}]{Leo Kozachkov}
\author[c]{Ksenia V. Kastanenka} 
\author[b]{Dmitry Krotov} 

\affil[a]{Department of Brain and Cognitive Sciences, MIT}
\affil[b]{MIT-IBM Watson AI Lab, IBM Research}
\affil[c]{Department of Neurology, MassGeneral Institute for Neurodegenerative Diseases, Massachusetts General Hospital and Harvard Medical School}


% Please give the surname of the lead author for the running footer
\leadauthor{Kozachkov} 


\maketitle

\begin{abstract}
Glial cells account for roughly 90\% of all human brain cells, and serve a variety of important developmental, structural, and metabolic functions. Recent experimental efforts suggest that astrocytes, a type of glial cell, are also directly involved in core cognitive processes such as learning and memory. While it is well-established that astrocytes and neurons are connected to one another in feedback loops across many timescales and spatial scales, there is a gap in our computational understanding of neuron-astrocyte interactions. To help bridge this gap, we draw on recent advances in artificial intelligence (AI) and astrocyte imaging technology. In particular, we show that neuron-astrocyte networks can naturally perform the core computation of a Transformer, a particularly successful type of AI architecture. In doing so, we provide a concrete and experimental testable account of neuron-astrocyte communication. Because Transformers are so successful across a wide variety of task domains, such as vision and audition, our analysis may help explain the ubiquity, flexibility, and power of the brain’s astrocyte-neuron networks.  
\end{abstract}


\begin{keywords}
Artificial Intelligence $|$ Neuroscience $|$ Transformers  $|$ Glia $|$ Astrocytes
\end{keywords}

\begin{corrauthor}
%\texttt{r.henriques{@}ucl.ac.uk}
leokoz8\at mit.edu, krotov\at ibm.com
\end{corrauthor}

\section*{Significance Statement}
Transformers have become the default choice of neural architecture for many machine learning applications. Their success across multiple domains such as language, vision, and speech raises the question: how can one build Transformers using biological computational units? At the same time, in the glial community there is a gradually accumulating evidence that astrocytes, formerly believed to be passive house-keeping cells in the brain, in fact play important role in brain's information processing and computation. In this work we hypothesize that astrocyte-neuron networks can naturally implement the core computation performed by the Transformer block in AI. The omnipresence of astrocytes in almost any brain area may explain the success of Transformers across a diverse set of information domains and computational tasks.

\section*{Introduction}

Astrocytes, one kind of glia, are a ubiquitous cell type in the central nervous system. There is increasing evidence that astrocytes play an active role in cognition \citep{kastanenka2020roadmap}. Specifically, astrocytes actively contribute to circuit activity underlying cognitive function. It is empirically well-established that astrocytes and neurons communicate with one another via feedback loops that span many spatial and temporal timescales \citep{Kofuji_Araque_2021,Lind_Brazhe_Jessen_Tan_Lauritzen_2013}. However, we lack a firm computational understanding of astrocyte-neuron communication.

Transformers, a particular type of AI architecture, have become influential in machine learning --and, increasingly, computational neuroscience--in the last few years \citep{Vaswani_Shazeer_Parmar_Uszkoreit_Jones_Gomez_Kaiser_Polosukhin_2017,toneva2019interpreting,krotov2020large,schrimpf2021neural,bricken2021attention,Whittington_Warren_Behrens_2022,caucheteux2022brains}. \lk{Describe transformers here}. They are currently the choice model for tasks across many disparate domains, including natural-language processing, vision, and speech \citep{lin2021survey}. Interestingly, several recent reports suggested representational similarities between Transformers and human brain recordings \citep{schrimpf2021neural,caucheteux2022brains,toneva2019interpreting}. However, unlike more traditional neural networks, such as feedforward convolutional neural networks \citep{Yamins_Hong_Cadieu_Solomon_Seibert_DiCarlo_2014} and Hopfield networks \citep{hopfield1982neural,krotov2020large}, Transformers lack a straightforward interpretation in terms of known biological processes. Thus, while there are tantalizing hints that Transformers and brains converge on similar solutions, we are lacking a concrete computational account for \textit{why}. 

Here we show that astrocyte-neuron networks perform the core computations of a Transformer. We arrive at this result by explicitly constructing an astrocyte-neuron network whose output, approximates that of a Transformer with high probability. The main novel computational element of our network is the tripartite synapse, the ubiquitous three-factor connection between an astrocyte, a pre-synaptic neuron, and a post-synaptic neuron \citep{Perea_Navarrete_Araque_2009}. We show that tripartite synapses perform the role of normalization in the Transformer’s self-attention operation. In the next section, we provide a brief primer on the biology of astrocytes and astrocyte-neuron interactions. 


Astrocytes are the other major cell type in the brain besides neurons. The exact ratio of glia to neurons is disputed, but it is somewhere between 1:1 and 10:1 \citep{von2016search} . The most well-studied type of glial cell is the astrocyte. A defining feature of astrocytes is that a single astrocyte cell forms connections with thousands to millions of nearby synapses \citep{halassa2007synaptic}. For example, a single human astrocyte can cover between 270,000 to 2 million synapses within a single domain 16. Astrocytes are for the most part electrically silent, encoding information in the dynamics of intracellular calcium ions ($\text{Ca}^{2+}$). In most parts of the brain, neurons and astrocytes are closely intertwined. For example, in the hippocampus as many as 60\% of all axon-dendrite synapses are wrapped by astrocyte cell membranes \citep{semyanov2021astrocytic}. In the cerebellum, the number is even higher. This three-way arrangement (pre-synaptic axon, post-synaptic dendrite, astrocytes membrane) is so common that it has been given a name: the tripartite synapse \citep{Perea_Navarrete_Araque_2009} . 

Astrocyte membranes contain receptors corresponding to the neurotransmitters released at the synaptic sites they ensheathe. For example, astrocytes in the basal ganglia are sensitive to dopamine, whereas in the cortex astrocytes are sensitive to glutamate. Despite being affected by the same pre-synaptic neurotransmitters, post-synaptic neurons and astrocytes respond very differently: neurons primarily encode information using action potentials, but astrocytes encode information via elevations in free intercellular calcium. Importantly, neuron-to-astrocyte signaling can trigger a response in the opposite astrocyte -to-neuron direction thus establishing a feedback loop between neural cells and astrocytes. Astrocytes can either depress or facilitate synapses, depending on the situation 18 . For example, astrocytes in the hypothalamus have been observed to multiplicatively scale the excitatory synapses they ensheathe by the same common factor \citep{Gordon_Iremonger_Kantevari_Ellis-Davies_MacVicar_Bains_2009} .

Interestingly, there is also extensive astrocyte-to- astrocyte communication in the brain. Astrocytes form large-scale networks with one another \citep{halassa2007synaptic}. These networks are spatially tiled, with regular intercellular spacing of a few tens of micrometers \citep{Sul_Orosz_Givens_Haydon_2004}.  Unlike neurons, who communicate primarily with spikes, astrocytes communicate via calcium waves that propagate between their cell bodies, processes, and endfeet \citep{Kuga_Sasaki_Takahara_Matsuki_Ikegaya_2011}. These waves have speeds of a few tens of micrometers per second. It is thought that these waves could be used to synchronize neural populations and coordinate important neural processes \citep{scemes2006astrocyte}.

\begin{figure*}[ht!]
\begin{center}
\includegraphics[width=2\columnwidth]{Figures/pnas_fig_1_long_version_2.png}
\caption{(Panel A) A high-level overview of the proposed neuron-astrocyte network. The Transformer block is approximated by a feed-forward network with an astrocyte unit that ensheaths the synapses between the hidden and last layers (matrix $H$). Data is constantly streamed into the network. (Panel B) During the writing phase the neuron-to-neuron weights are updated using Hebbian learning rule and the neuron-to-astrocyte are updated using a presynaptic plasticity rule . During the reading phase the data is forwarded through the network, and the astrocyte modulates the synaptic weights $H$.}
\label{fig:macro_bio_drawing}
\end{center}
\end{figure*}


\section*{Biological Implementation of a Transformer Block}
This section begins by introducing Transformers using AI notations. Then, the neuron-astrocyte network is described, and it is shown that the attention operation can be implemented using the proposed model. At the end of the section it is explained how the remaining operations of the Transformer block (layer normalization and feed-forward network) can be included into the neuron-astrocyte circuit.  
\subsection{Transformer Notation}
Consider a sequence of $N$ token embeddings. Each token can correspond to a word (or a part of the word) if the Transformer is used in the language domain, or a patch of an image in the vision domain. Each embedding is of dimension $\edim$. The tokens are streamed into the network one-by-one (online setting), and the time of the token's presentation is denoted by $t$. The $t^{th}$ embedding is given by a vector $\bx_t \in \fR^\edim$. In the Transformer block each of these tokens is converted to a key, value, or query vector via a corresponding linear transformation, $W_K,W_Q \in \fR^{D \times \edim}$ and $W_V \in \fR^{d \times d}$:
\begin{align}\label{eq:transformer_kvq}
\begin{split}
\bk_t = W_K \bx_t\\
\bv_t = W_V \bx_t \\
\bq_t = W_Q \bx_t\\
\end{split}
\end{align}
These vectors are collected into matrices $X,K,V,Q$ by stacking them columwise. The nonlinearities are applied to matrices columnwise:
\begin{align*}
\phi(K) \equiv \begin{bmatrix}
| & | & & |\\
\phi(\bk_1) & \phi(\bk_2) & \dots & \phi(\bk_N) \\
| & | & & | 
\end{bmatrix}\
\end{align*}
Written in this way, the Transformer self-attention operator becomes:

\begin{equation}\label{eq:self-attention}
\text{SelfAttn}(X) = V\text{softmax}(K^TQ)
\end{equation}
where the softmax normalization is computed along the \textit{columns} of $K^TQ$. The output of this self-attention operation is then passed along to a feed-forward neural network (FFN) that acts separately on each token (each column of its input). Without loss of generality, a single-headed attention Transformer is studied. In this case the output of the full Transformer block may be written as a two-step process:
\begin{align}\label{eq:Transformer_Block}
\begin{split}
Y = \text{LayerNorm}(\text{SelfAttn}(X) + X) \\
\text{Transformer}(X) = \text{LayerNorm}(\text{FFN}(Y) + Y)
\end{split}
\end{align}
where FFN refers to a feedfoward network, applied to each token separately and identically. It is not obvious how or if it is possible to write \eqref{eq:Transformer_Block} in terms of simple mathematically abstracted biological operations (artificial neurons and perhaps astrocytes). Note that the main difficulty is that the $\text{SelfAttn}$ operation contains interaction terms $between$ token embeddings \citep{Lu_Li_He_Sun_Dong_Qin_Wang_Liu_2019}, and not just between elements of a single embedded token. In the next few sections, we show that by defining Hebbian-based ``read'' and ``write'' operations, we can capture these interactions in a biologically plausible way.  

In order to gain theoretical insight into Transformers, it is common to tie the weights \citep{sander2022sinkformers,yang2022transformers}. This tying can be within a single Transformer block, between blocks, or both. Going forward we will tie the weights within a single block, but not between blocks. In particular, we tie $W_Q,W_K,W_V$ as follows:
\[W_Q = W_K = \biow, \hspace{.2cm} W_V = I \]
for some arbitrary matrix $\biow$ and the identity matrix, $I$. In general, we will not require that $d = D$. We include this constraint now to fully analyze the simplest version of our model that captures the essential elements of our argument. Later on in section \ref{section: untied_weights} we extend to more specialized cases where the weights are not tied. Without loss of generality, we will ignore layer normalization steps for now, returning to it in section \ref{section: normalization}.


\subsection{Neuron-Astrocyte Network}
A high-level overview of our circuit is shown in Figure \ref{fig:macro_bio_drawing}. The  network consists of a perceptron with an input layer, a hidden layer, and an output layer. The $\edim$-dimensional inputs are passed to the hidden layer with $m$ units, as well as to the last layer via a skip connection (not shown). The hidden layer applies a fixed nonlinearity to incoming inputs. The outputs of the hidden layer are passed to the last layer via a linear mapping $H \in \fR^{d \times m}$. The synapses in the matrix $H$ are triparite synapses, meaning that each of the $m\edim$ synapses is associated with an astrocyte process $p_{i \alpha}$. The Latin indices $i,j$ are used to enumerate neurons in the first and last layers, while the Greek indices $\alpha, \beta$ are reserved for the hidden neurons. With these notations the strength of the synapse between a hidden neuron $\alpha$ and the output neuron $i$ is denoted by $H_{i\alpha}$ and the activity of the astrocyte process that ensheaths this synapse is described by $p_{i \alpha}$. The layers are denoted from left to right as $\beff$,$\bh$,$\bell$ (first, hidden, last), respectively. Our network is described by the following equations:
\begin{align}\label{eq:foward_eqs}
\begin{split}
\beff = \bx & \hspace{.5cm} \in \fR^{d} \\ 
\bh = \phi(\biow\beff) & \hspace{.5cm} \in \fR^{m} \\
\bell = r(H\odot\tilde{P})\bh + \beff   & \hspace{.5cm} \in \fR^{d} \\    
\end{split}
\end{align}
The scalar $r = \{0,1\}$ stands for `read', and is zero during the writing phase and unity during the reading phase. The symbol $\odot$ denotes the Hadamard product (element-wise multiplication) between two matrices. The matrix $\tilde{P} \in \fR^{\edim \times m}$ captures the effect of the astrocyte processes, and is defined as follows:
\[\tilde{P}_{i \alpha} = \frac{1}{p_{i \alpha}}\]
This inverse modulation of synaptic weights by astrocytes has been observed, for example, in studies involving tumour necrosis factor-alpha (TNF-$\alpha$), wherein astrocytes will upscale synaptic weights in response to low neural activity and downscale weights in response to high neural activity. More generally, many studies have observed that astrocytes can both depress and facilitate synapses, depending on the situation \cite{perea2007astrocytes, perea2009tripartite, de2016astrocytes,Kofuji_Araque_2021,ivanov2021increasing}


\subsection{Astrocyte Process Dynamics}
As discussed in the introduction, astrocyte processes are sensitive to pre-synaptic neural activity. To capture this mathematically, we assume that the astrocyte process $\ca$ response is linearly proportional to the pre-synaptic neuron activation $h_\alpha$ of neuron $\alpha$ in layer $\bh$. The constant of proportionality between the astrocyte process activation and the pre-synaptic neural activity is denoted as $g_{i \alpha}$. This constant is in general different for every astrocyte process. Upon presentation of an embedded token to the network, astrocyte process $p_{i \alpha}$ initially responds with a local calcium elevation $g_{i \alpha} h_{\alpha}$. This $\ca$ response is then spatially averaged with the responses of other nearby astrocyte processes so that, after transients, the processes have the same value once a token is presented:

\begin{align}\label{eq:astrocyte_process}
\begin{split}
p_{i \alpha} = \frac{1}{md}\sum^{\edim}_{j = 1} \sum^{m}_{\beta = 1}g_{j \beta}h_\beta
\end{split}
\end{align}
The neuron-to-astrocyte signalling pathway in our circuit is completely described by equation \eqref{eq:astrocyte_process}.


\subsection{Writing Phase}
During the writing phase, $r$ is set to zero. Biologically, this condition could correspond to some global neuromodulator being released into the local environment, for example acetylcholine, as suggested in \citep{rasmusson2000role,Tyulmankov_Fang_Vadaparty_Yang_2021}. Plugging $r = 0$, \eqref{eq:foward_eqs} becomes: 

\begin{align}\label{eq:foward_eqs_write}
\begin{split}
\beff_t = \bx_t\\ 
\bh_t = \phi(\biow\beff_t) = \phi(\bk_t)\\
\bell_t = \beff_t = \bv_t\\     
\end{split}
\end{align}
As the embedded tokens are passed into \eqref{eq:foward_eqs_write} sequentially, the weight matrix $H$ is updated via Hebbian plasticity with a learning rate of $\frac{1}{m}$. Upon presentation of token $t$, the matrix $H$ is:

\[H_t = H_{t-1} +  \frac{1}{m}\bell_t \bh_t^T =  H_{t-1} + \frac{1}{m}\bv_t\phi^T(\bk_t) \]
After all the embeddings are passed through the network, assuming $H$ is initially the zero matrix, the final value of $H$ is:

\[H= \frac{1}{m}V\phi^T(K) \]
At the same time, the neuron-to-astrocyte weights are updated via pre-synaptic plasticity. Upon presentation of token $t$, these weights are: 

\[g_{t, i \alpha} = g_{t-1, i \alpha}+ \phi(\biow\bx_t)_\alpha \rightarrow g_{i \alpha} = \sum^{N}_{t = 1}\phi(\bk_t)_{\alpha}\]
Note that as a consequence of the pre-synaptic plasticity, the weight $g_{i \alpha}$ does not depend on the index $i$. Therefore, we will only refer to the vector $\bg \in \fR^{m}$, which--through the pre-synaptic plasticity--is simply the sum over all token presentations of the hidden layer neural activations:

\[\bg = \sum^{N}_{t = 1}\phi(\bk_t)\]

\subsection{Reading Phase}
During the reading phase, the read gate is set to $r = 1$ in \eqref{eq:foward_eqs}, and the inputs are forwarded through the network. The astrocyte process activation value $p$, which according to \eqref{eq:astrocyte_process} does not depend on indices $i$ and $\alpha$, is given by:
\begin{align}
\begin{split}
p = \frac{1}{m}\bh^T\bg = \frac{1}{m}\phi(\bq_t)^T\sum^{N}_{t = 1}\phi(\bk_t) 
\end{split}    
\end{align}
To obtain the last equality we have used $\bh_t = \phi(\biow\bx_t) = \phi(\bq_t)$. Plugging in all the steps of \eqref{eq:foward_eqs}, we see that the last layer has the following value:

\begin{equation}\label{eq:token_wise_approx}
\bell_t = \frac{1}{p}H\phi(\bq_t) + \bx_t =  \frac{V\phi^T(K)\phi(\bq_t)}{\phi(\bq_t)^T\sum^{N}_{t = 1}\phi(\bk_t)} + \bx_t     
\end{equation}
If we compute $\bell$ for every input $\bx_t$ and stack the results column-wise into a matrix $L$, we have:
\begin{equation}\label{eq:bio_out}
L = V\phi^T(K)\phi(Q)G^{-1} + X
\end{equation}
where $G$ is a diagonal matrix such that $G_{zz} = \phi(\bq_z)^T\sum^{N}_{t = 1}\phi(\bk_t)$. 

\begin{figure}[ht]
\begin{center}
\includegraphics[width=1.0\linewidth]{Figures/error_plot.png}
\caption{Error vs number of hidden units ($m$) in our network. As $m$ increases, the difference between the output of the biological circuit and the `pure' Transformer block decreases.}
\label{fig:selfattnvsbio}
\end{center}
\end{figure}

At this point, \eqref{eq:bio_out} is equivalent to a non-autoregressive version of the Linear Transformer \citep{Katharopoulos_Vyas_Pappas_Fleuret_2020}. However, to make our circuit approximate a traditional Transformer with softmax attention, one more piece is required. We need to choose $\phi$ carefully so that we can reproduce the exponential dot-product in the softmax operation. In other words, we need $\phi$ to satisfy: 

\[ \phi(\bx)^T\phi(\mathbf{y}) \approx \text{exp}(\bx^T\mathbf{y}) \]
This can be achieved using random feature attention (RFA) \citep{peng2021random}. RFA relies on a well-known result in kernel approximation theory, which is that the radial basis function (RBF) kernel can, with high probability, be approximated very well using random projections and cosines \citep{rahimi2007random}. To map the RBF kernel to the exponential dot-product kernel used in Transformers, one can simply scale the RBF approximation as follows:

\begin{equation} \label{eq:phi_rbf_approx}
\phi(\bx) = \text{exp}(\frac{||\bx||^2}{2})\phi_{\text{RBF}}(\bx) = \text{exp}(\frac{||\bx||^2}{2})\text{cos}(\Pi\bx + \bb)  
\end{equation}
where the elements of $\Pi \in \fR^{m \times D}$ are drawn from a standard normal distribution, and the elements of $\bb \in \fR^{m}$ are drawn from the uniform distribution on $[0,2\pi]$. The cosine is applied element-wise. Note that we have dropped the usual factor of $\sqrt{\frac{2}{m}}$ from the RBF approximation of  \cite{rahimi2007random}--this is because all constant prefactors of $\phi$ get canceled out in the fraction appearing in \eqref{eq:token_wise_approx}. Plugging \eqref{eq:phi_rbf_approx} into \eqref{eq:bio_out} yields the following:


\begin{align}\label{eq:bio_out_approx}
\begin{split}
L \approx \text{SelfAttn}(X) + X
\end{split}
\end{align}


\subsection{The Role of Normalization and Feedforward Layers}\label{section: normalization}
Thus far we have neglected the role of layer normalization and feedfoward layers. That is because these are operations that are performed on a \textit{per-token} basis, and can therefore be dropped into our model without changing anything in our arguments. In the Transformer model, layer normalization occurs directly preceding and succeeding the FFN layer. With layer normalization and FFN layer, our circuit becomes:

\begin{align*}
\begin{split}
Y = \text{LayerNorm}(L) \approx \text{LayerNorm}(\text{SelfAttn}(X) + X)  \\
\text{Transformer}(X) = \text{LayerNorm}(\text{FFN}(Y) + Y)
\end{split}
\end{align*}
Note that both the FFN and layer norm operations are biologically plausible. Layer normalization is a form of divisive normalization, which is well-studied in the neuroscience literature \citep{carandini2012normalization,Shen_Wang_Navlakha_2021}. In the context of Transformers, it can be achieved by inserting a hidden layer (with lateral connections to implement the normalization) directly before and after the feedforward layer.

\section*{General Case of Untied Weights}\label{section: untied_weights}
In this section we relax the weight tying condition, and generalize our construction to the case when $D \neq d$. Whereas in the previous sections $r$ acted as a gatekeeper for weight matrix $H$, we will now \textit{also} have $r$ act a gatekeeper for a few other weight matrices. Using the same variable names, consider the following neuron-astrocyte forward equations:

\begin{align}\label{eq:foward_eqs_general}
\begin{split}
\beff = \bx & \hspace{.5cm} \in \fR^{d} \\ 
\bh = (1-r)\phi(\biow_K\beff) + r\phi(\biow_Q\beff) & \hspace{.5cm} \in \fR^{m} \\
\bell = r(H\odot\tilde{P})\bh +  (1-r)\biow_V\beff + r\beff    & \hspace{.5cm} \in \fR^{d} \\    
\end{split}
\end{align}
When $r = 0$, we recover the writing phase of \eqref{eq:foward_eqs_write}; when $r=1$, we recover  the reading phase equations of \eqref{eq:token_wise_approx}. When we impose the weight tying constraint of $W_K = W_Q = W$ and $W_V = I$, we recover the original equations of \eqref{eq:foward_eqs}.

\section*{Numerical Validation} The results derived above have also been checked numerically. In Figure \ref{fig:selfattnvsbio} one can see the error between the proposed neuron-astrocyte network and the Transformer block as a function of the ratio of the width of the hidden layer to the size of the token embedding. As expected from the theoretical analysis, the error between the two networks rapidly decreases as the hidden layer becomes wider. In practice, as the width of the hidden layer becomes 5-10 times the embedding dimension the two networks produce very similar outputs.  

\section*{Do We Need Astrocytes?}
Although we are interested in relating Transformers to glial cells, a natural question when positing any new brain mechanism is: ``Can the same behavior be achieved with the old, well-known mechanisms?''. This section demonstrates that an equivalent Transformer circuit can be constructed using just neurons, together with divisive normalization achieved via shunting inhibition. The circuit is similar to \eqref{eq:foward_eqs}:

\begin{align}\label{eq:inhibitory_circuit}
\begin{split}
\beff = \bx & \hspace{.5cm} \in \fR^{d} \\ 
\bh = \phi(\biow\beff) & \hspace{.5cm} \in \fR^{m} \\
R = \bg^T \bh & \hspace{.5cm} \in \fR \\
\bell = \frac{r}{R} H\bh + \beff   & \hspace{.5cm} \in \fR^{d} 
\end{split}
\end{align}
The only difference between \eqref{eq:inhibitory_circuit} and \eqref{eq:foward_eqs} is the addition of a new element, $R$, and the removal of the astrocyte processes. Here $R$ is an inhibitory neuron that divisively normalizes feedfoward inputs into layer $\bell$. However, it does not inhibit all feedfoward inputs equally. Despite both $\bh$ and $\beff$ being feedfoward inputs to layer $\bell$, the divisive inhibition is only implemented on the inputs coming from layer $\bh$. This can happen, for example, if the feedfoward synaptic inputs coming from layer $\bh$ arrive at the dendritic tree close to where inhibitory inputs from neuron $R$ shunt current flow, while the feedfoward inputs coming from layer $\beff$ synapse far away from the shunting \citep{chance2000divisive}. Leaving the reading and writing phases untouched, circuit \eqref{eq:inhibitory_circuit} implements the same forward pass as \eqref{eq:foward_eqs}.


\section*{Time Scale of Transformers in the Brain}
A notable component explicitly missing from Transformers, as well as from our biological circuit, is time. Unlike the brain, Transformers are not dynamical systems. Except in the special instance of autoregressive linear Transformers (which may be seen as a kind of recurrent neural network), Transformers do not maintain a hidden state via recurrent weights. Nevertheless, timing is a critical aspect of information processing in the brain, so in this section we describe a plausible timescale for our circuit in terms of known biophysics parameters. The following paragraph argues that the total compute time for our circuit is on the order of $(N+1) \times 100$ milliseconds, where $N$ is the number of tokens. 
Our circuit operates in two distinct phases: a reading phase and a writing phase. The reading phase does not involve any plasticity, so the only relevant timescale to compute is how long it takes to traverse the neuron $\rightarrow$ astrocyte $\rightarrow$ synapse pathway. Recent data indicate that this entire signaling pathway can be traversed on the order of a few hundreds of milliseconds \citep{stobart2018cortical}. The speed of the writing phase is limited by the speed of plasticity. There are two types of plasticity used in our model: 1) Hebbian plasticity between neurons and 2) pre-synaptic plasticity between neuronal and astrocytic processes. The former can happen on relatively fast timescale, for example through spike-timing-dependent plasticity (STDP). In that case of STDP, the timescale can be as fast as a few tens of milliseconds 23 . The latter timescale is harder to establish, due to limitations in calcium recording technology. While fast calcium transients in astrocyte processes have been recently recorded \citep{stobart2018cortical}, and neuron-astrocyte plasticity has been experimentally observed \citep{croft2015plasticity} , fast (e.g < 1s) neuron-astrocyte plasticity has not been observed. Assuming the timescale of this plasticity is also a few hundred milliseconds, then an estimate for the overall timescale of our model would be a few hundred milliseconds for the read phase, plus a few hundred milliseconds per token for the write phase. This leads to a total compute time of $\tau_{compute} = (100 + N \times 100)$ milliseconds. Finally, note that most "real-world" Transformers are composed of many blocks. Our circuit is a model of a single block. Thus, the total compute time for a $k$ block transformer modelled using our framework is $k\tau_{compute}$.


\section*{Discussion}
Here we have built a computational astrocyte-neuron model which is functionally equivalent to an important AI architecture: the Transformer. This model serves a dual purpose. The first purpose is to provide a concrete, computational account of how the communication between astrocytes and neurons subserves brain function. The second purpose is to provide a biologically-plausible account of how Transformers might be implemented in the brain.
While the feedback loop between neurons and astrocytes is well-studied from an experimental perspective, there is comparatively little work studying it from a computational perspective 1. Most of the computational studies (with notable exceptions \citep{song2016design,ivanov2021increasing,tang2019introducing}) are focused on understanding the biophysics of neuron-astrocyte or astrocyte-astrocyte signaling. An important difference between our model and previous models of astrocyte-neuron interactions is that our model is generic, that is any Transformer is an astrocyte-neuron network. Given the demonstrated power and flexibility of Transformers, this generality can help to explain why astrocytes are so prevalent across disparate brain areas and species. 
Our model has several immediate applications. First, as calcium imaging technologies improve, it will become increasingly feasible to explicitly compare artificial representations in AI networks to representations in biological astrocyte networks—as is already done when comparing AI networks to biological neural networks \citep{schrimpf2021neural,Yamins_Hong_Cadieu_Solomon_Seibert_DiCarlo_2014,schrimpf2020brain}. Our work suggests that a reasonable place to start may be to compare Transformer normalization terms to astrocytic calcium responses. Second, our work suggests that experimenters may perform causal manipulations on Transformers to generate putative hypotheses for how astrocyte function goes astray in brain disorders and diseases \citep{escartin2021reactive}. These hypotheses can in turn be potentially used to suggest novel treatments and routes for clinical success.


\begin{acknowledgements}
This work was completed during the MIT-IBM Watson AI Lab Summer 2022 Internship Program. We thank John Hopfield and Mriganka Sur for helpful comments and feedback. 
\end{acknowledgements}

\section*{Bibliography}
\bibliography{bio_transformer}

\end{document}
