## Supplementary material for "Building Transformers from Neurons and Astrocytes": LaTeX files: main_10042022.tex

\RequirePackage[2020-02-02]{latexrelease}
\documentclass[times, twoside]{zHenriquesLab-StyleBioRxiv}
\usepackage{blindtext}
\usepackage{amsmath}
\usepackage{authblk}
\usepackage{amsfonts}
\usepackage{mathtools}
\usepackage{accents}
\usepackage{graphicx}
\usepackage{amssymb}
\usepackage{xcolor}

\newcommand{\lk}[1]{\textcolor{red}{Leo: #1}}
\newcommand{\dk}[1]{\textcolor{blue}{Dima: #1}}
\input{preamble}

% Please give the surname of the lead author for the running footer
\leadauthor{Henriques} 

\begin{document}

\title{Building Transformers from Neurons and Glia}
\shorttitle{Building Transformers from Neurons and Glia}

% Use letters for affiliations, numbers to show equal authorship (if applicable) and to indicate the corresponding author
\author[a,b,\footnote{Work completed during MIT-IBM Watson AI Lab Summer 2022 Internship.}]{Leo Kozachkov}
\author[c]{Ksenia V. Kastanenka} 
\author[b]{Dmitry Krotov} 

\affil[a]{Department of Brain and Cognitive Sciences, MIT}
\affil[b]{MIT-IBM Watson AI Lab, IBM Research}
\affil[c]{Department of Neurology, MassGeneral Institute for Neurodegenerative Diseases, Massachusetts General Hospital and Harvard Medical School}


% Please give the surname of the lead author for the running footer
\leadauthor{Kozachkov} 


\maketitle

\begin{abstract}
We argue that neuron-astrocyte networks can naturally perform the computation of a Transformer. The cornerstone of the argument is a simple mathematical model of a neuron-astrocyte network that approximates the forward pass of a Transformer block to arbitrary accuracy. The main idea behind our approach is the shift from a neuron-centric design to a hybrid neuron-glia architecture with direct synaptic modulation by astrocyte cells. The main computational element of our model is a mathematical abstraction of a tripartite synapse—the ubiquitous three-factor connection between pre- and post-synaptic neurons, as well as the astrocyte cell. The circuit alternates between periods of writing and reading. During the writing period, inputs are stored via Hebbian learning. During the reading period, inputs are simply passed through the network. The model relies on plasticity between the pre- and post-synaptic neurons, plasticity between the pre-synaptic neuron and the astrocyte, and the astrocyte-dependent modulation of the synapses between the neurons. We argue that there exists substantial experimental evidence for all these phenomena. As such, neuron-astrocyte networks are natural candidates for the biological ``hardware'' that can be used for conducting computation with Transformers.
\end{abstract}


\begin{keywords}
Artificial Intelligence $|$ Neuroscience $|$ Transformers  $|$ Glia $|$ Astrocytes
\end{keywords}

\begin{corrauthor}
%\texttt{r.henriques{@}ucl.ac.uk}
leokoz8\at mit.edu, krotov\at ibm.com
\end{corrauthor}

\section*{Significance Statement}
Transformers have become the default choice of neural architecture for many machine learning applications. Their success across multiple domains such as language, vision, and speech raises the question: how can one build Transformers using biological computational units? At the same time, in the glial community there is a gradually accumulating evidence that astrocytes, formerly believed to be passive house-keeping cells in the brain, in fact play important role in brain's information processing and computation. In this work we hypothesize that astrocyte-neuron networks can naturally implement the core computation performed by the Transformer block in AI. The omnipresence of astrocytes in almost any brain area may explain the success of Transformers across a diverse set of information domains and computational tasks.

\section*{Introduction}
Transformers \citep{Vaswani_Shazeer_Parmar_Uszkoreit_Jones_Gomez_Kaiser_Polosukhin_2017} have become extremely influential in the last few years. They are currently the choice model for tasks across many disparate domains, including natural-language processing, vision, and speech \citep{lin2021survey}. Because of Transformer's success, many researchers are exploring their connections to biological computations in the brain \citep{toneva2019interpreting,krotov2020large,schrimpf2021neural,bricken2021attention,Whittington_Warren_Behrens_2022,caucheteux2022brains}. A key question for this line of research is: 

\textit{Can a Transformer be built using simple, biologically-plausible operations}? 

Here we argue that the answer is: \textit{yes}. We arrive at this result by explicitly constructing a biological circuit whose output can--with high probability--approximate that of a Transformer's to an arbitrary degree of accuracy. The core computational element of our network is a ``three-factor'' synapse, which fulfils the role of normalization in the Transformer's self-attention operation. In the sections below we argue that the function of these three-factor connections could be achieved in the brain by astrocytes, a type of glial cell that wrap and modulate nearby synapses to from tripartite synapses \citep{kandel2000principles,kastanenka2020roadmap}.


Astrocytes, whose role was long thought to be neural ``housekeeping'', have recently emerged as potential key players in brain computation \citep{kastanenka2020roadmap,Kofuji_Araque_2021}. Despite this, few hypotheses have been put forward regarding the specific role of astrocytes in brain computation. Our aim in this paper is to help close this knowledge gap. We do so by establishing a one-to-one relationship between Transformers and biological neuron-astrocyte networks, thereby providing a specific computational hypothesis for how the dialog between neurons and astrocytes subserves cognition. 

Glial cells are the other major cell type in the brain besides neurons. The exact ratio  of glia to neurons is disputed, but it is somewhere between 1:1 and 10:1 \citep{von2016search}. Glial cells are for the most part electrically passive, encoding information in the dynamics of intracellular calcium ions ($\text{Ca}^{2+}$). In most parts of the brain, there is a close apposition of neurons and glial cells. For example, in the hippocampus as many as 60\% of all axon-dendrite synapses are wrapped by glial cell membranes \citep{verkhratsky2007glial}. In the cerebellum, the number is even higher. This three-way arrangement (pre-synaptic axon, post-synaptic dendrite, glial membrane) is so common that it has been given a name: \textit{the tripartite synapse} \citep{Perea_Navarrete_Araque_2009}. 

The most well-studied type of glial cell is the astrocyte. A defining feature of astrocytes is that a single astrocyte cell forms tripartite synapses with thousands to millions of nearby synapses \citep{halassa2007synaptic}. For example, a single human astrocyte can cover between 270,000 to 2 million synapses within a single domain \citep{Oberheim_2009}. Glial membranes tend to be have receptors corresponding to the neurotransmitters released at the synaptic site they ensheath. For example, astrocytes in the basal ganglia are sensitive to dopamine, whereas in the cortex astrocytes are sensitive to glutamate. Despite being affected by the same pre-synaptic neurotransmitters, post-synaptic neurons and glia respond very differently: neurons will respond with a post-synaptic action potential, but glia will respond with an elevation in free intercellular $\text{Ca}^{2+}$. Importantly, neuron-to-glia signalling can trigger a reaction in the other direction (glia-to-neuron) thus establishing a feedback loop between neural cells and glial cells. Glia can either depress or facilitate synapses, depending on the situation \citep{Newman_2003}. For example, astrocytes in the hypothalamus have been observed to multiplcatively scale the excitatory synapses they ensheath by the same common factor \citep{Gordon_Iremonger_Kantevari_Ellis-Davies_MacVicar_Bains_2009}.

Interestingly, there is also extensive glial-to-glial communication in the brain. For example, astrocytes form large-scale networks with one another \citep{halassa2007synaptic}. These networks are spatially tiled, with regular intercellular spacing of a few tens of micrometers \citep{Sul_Orosz_Givens_Haydon_2004}.  Unlike neurons, who communicate with spikes, astrocytes communicate via calcium waves that propagate between their cell bodies \citep{Kuga_Sasaki_Takahara_Matsuki_Ikegaya_2011}. These waves have speeds of a few tens of micrometers per second. It is thought that these waves could be use to synchronize neural populations and coordinate important neural processes \citep{scemes2006astrocyte}.
\begin{figure*}[ht!]
\begin{center}
\includegraphics[width=2\columnwidth]{Figures/pnas_fig_1_long_version_2.png}
\caption{(Panel A) A high-level overview of the proposed neuron-astrocyte network. The Transformer block is approximated by a feed-forward network with an astrocyte unit that ensheaths the synapses between the hidden and last layers (matrix $H$). Data is constantly streamed into the network. (Panel B) During the writing phase the neuron-to-neuron weights are updated using Hebbian learning rule and the neuron-to-astrocyte are updated using a presynaptic plasticity rule . During the reading phase the data is forwarded through the network, and the astrocyte modulates the synaptic weights $H$.}
\label{fig:macro_bio_drawing}
\end{center}
\end{figure*}


\section*{Biological Implementation of a Transformer Block}
This section begins by introducing Transformers using AI notations. Then, the neuron-astrocyte network is described, and it is shown that the attention operation can be implemented using the proposed model. At the end of the section it is explained how the remaining operations of the Transformer block (layer normalization and feed-forward network) can be included into the neuron-astrocyte circuit.  
\subsection{Transformer Notation}
Consider a sequence of $N$ token embeddings. Each token can correspond to a word (or a part of the word) if the Transformer is used in the language domain, or a patch of an image in the vision domain. Each embedding is of dimension $\edim$. The tokens are streamed into the network one-by-one (online setting), and the time of the token's presentation is denoted by $t$. The $t^{th}$ embedding is given by a vector $\bx_t \in \fR^\edim$. In the Transformer block each of these tokens is converted to a key, value, or query vector via a corresponding linear transformation, $W_K,W_Q \in \fR^{D \times \edim}$ and $W_V \in \fR^{d \times d}$:
\begin{align}\label{eq:transformer_kvq}
\begin{split}
\bk_t = W_K \bx_t\\
\bv_t = W_V \bx_t \\
\bq_t = W_Q \bx_t\\
\end{split}
\end{align}
These vectors are collected into matrices $X,K,V,Q$ by stacking them columwise. The nonlinearities are applied to matrices columnwise:
\begin{align*}
\phi(K) \equiv \begin{bmatrix}
| & | & & |\\
\phi(\bk_1) & \phi(\bk_2) & \dots & \phi(\bk_N) \\
| & | & & | 
\end{bmatrix}\
\end{align*}
Written in this way, the Transformer self-attention operator becomes:

\begin{equation}\label{eq:self-attention}
\text{SelfAttn}(X) = V\text{softmax}(K^TQ)
\end{equation}
where the softmax normalization is computed along the \textit{columns} of $K^TQ$. The output of this self-attention operation is then passed along to a feed-forward neural network (FFN) that acts separately on each token (each column of its input). Without loss of generality, a single-headed attention Transformer is studied. In this case the output of the full Transformer block may be written as a two-step process:
\begin{align}\label{eq:Transformer_Block}
\begin{split}
Y = \text{LayerNorm}(\text{SelfAttn}(X) + X) \\
\text{Transformer}(X) = \text{LayerNorm}(\text{FFN}(Y) + Y)
\end{split}
\end{align}
where FFN refers to a feedfoward network, applied to each token separately and identically. It is not obvious how or if it is possible to write \eqref{eq:Transformer_Block} in terms of simple mathematically abstracted biological operations (artificial neurons and perhaps astrocytes). Note that the main difficulty is that the $\text{SelfAttn}$ operation contains interaction terms $between$ token embeddings \citep{Lu_Li_He_Sun_Dong_Qin_Wang_Liu_2019}, and not just between elements of a single embedded token. In the next few sections, we show that by defining Hebbian-based ``read'' and ``write'' operations, we can capture these interactions in a biologically plausible way.  

In order to gain theoretical insight into Transformers, it is common to tie the weights \citep{sander2022sinkformers,yang2022transformers}. This tying can be within a single Transformer block, between blocks, or both. Going forward we will tie the weights within a single block, but not between blocks. In particular, we tie $W_Q,W_K,W_V$ as follows:
\[W_Q = W_K = \biow, \hspace{.2cm} W_V = I \]
for some arbitrary matrix $\biow$ and the identity matrix, $I$. In general, we will not require that $d = D$. We include this constraint now to fully analyze the simplest version of our model that captures the essential elements of our argument. Later on in section \ref{section: untied_weights} we extend to more specialized cases where the weights are not tied. Without loss of generality, we will ignore layer normalization steps for now, returning to it in section \ref{section: normalization}.


\subsection{Neuron-Astrocyte Network}
A high-level overview of our circuit is shown in Figure \ref{fig:macro_bio_drawing}. The  network consists of a perceptron with an input layer, a hidden layer, and an output layer. The $\edim$-dimensional inputs are passed to the hidden layer with $m$ units, as well as to the last layer via a skip connection (not shown). The hidden layer applies a fixed nonlinearity to incoming inputs. The outputs of the hidden layer are passed to the last layer via a linear mapping $H \in \fR^{d \times m}$. The synapses in the matrix $H$ are triparite synapses, meaning that each of the $m\edim$ synapses is associated with an astrocyte process $p_{i \alpha}$. The Latin indices $i,j$ are used to enumerate neurons in the first and last layers, while the Greek indices $\alpha, \beta$ are reserved for the hidden neurons. With these notations the strength of the synapse between a hidden neuron $\alpha$ and the output neuron $i$ is denoted by $H_{i\alpha}$ and the activity of the astrocyte process that ensheaths this synapse is described by $p_{i \alpha}$. The layers are denoted from left to right as $\beff$,$\bh$,$\bell$ (first, hidden, last), respectively. Our network is described by the following equations:
\begin{align}\label{eq:foward_eqs}
\begin{split}
\beff = \bx & \hspace{.5cm} \in \fR^{d} \\ 
\bh = \phi(\biow\beff) & \hspace{.5cm} \in \fR^{m} \\
\bell = r(H\odot\tilde{P})\bh + \beff   & \hspace{.5cm} \in \fR^{d} \\    
\end{split}
\end{align}
The scalar $r = \{0,1\}$ stands for `read', and is zero during the writing phase and unity during the reading phase. The symbol $\odot$ denotes the Hadamard product (element-wise multiplication) between two matrices. The matrix $\tilde{P} \in \fR^{\edim \times m}$ captures the effect of the astrocyte processes, and is defined as follows:
\[\tilde{P}_{i \alpha} = \frac{1}{p_{i \alpha}}\]
This inverse modulation of synaptic weights by astrocytes has been observed, for example, in studies involving tumour necrosis factor-alpha (TNF-$\alpha$), wherein astrocytes will upscale synaptic weights in response to low neural activity and downscale weights in response to high neural activity. More generally, many studies have observed that astrocytes can both depress and facilitate synapses, depending on the situation \cite{perea2007astrocytes, perea2009tripartite, de2016astrocytes,Kofuji_Araque_2021,ivanov2021increasing}


\subsection{Astrocyte Process Dynamics}
As discussed in the introduction, astrocyte processes are sensitive to pre-synaptic neural activity. To capture this mathematically, we assume that the astrocyte process $\ca$ response is linearly proportional to the pre-synaptic neuron activation $h_\alpha$ of neuron $\alpha$ in layer $\bh$. The constant of proportionality between the astrocyte process activation and the pre-synaptic neural activity is denoted as $g_{i \alpha}$. This constant is in general different for every astrocyte process. Upon presentation of an embedded token to the network, astrocyte process $p_{i \alpha}$ initially responds with a local calcium elevation $g_{i \alpha} h_{\alpha}$. This $\ca$ response is then spatially averaged with the responses of other nearby astrocyte processes so that, after transients, the processes have the same value once a token is presented:

\begin{align}\label{eq:astrocyte_process}
\begin{split}
p_{i \alpha} = \frac{1}{md}\sum^{\edim}_{j = 1} \sum^{m}_{\beta = 1}g_{j \beta}h_\beta
\end{split}
\end{align}
The neuron-to-astrocyte signalling pathway in our circuit is completely described by equation \eqref{eq:astrocyte_process}.


\subsection{Writing Phase}
During the writing phase, $r$ is set to zero. Biologically, this condition could correspond to some global neuromodulator being released into the local environment, for example acetylcholine, as suggested in \citep{rasmusson2000role,Tyulmankov_Fang_Vadaparty_Yang_2021}. Plugging $r = 0$, \eqref{eq:foward_eqs} becomes: 

\begin{align}\label{eq:foward_eqs_write}
\begin{split}
\beff_t = \bx_t\\ 
\bh_t = \phi(\biow\beff_t) = \phi(\bk_t)\\
\bell_t = \beff_t = \bv_t\\     
\end{split}
\end{align}
As the embedded tokens are passed into \eqref{eq:foward_eqs_write} sequentially, the weight matrix $H$ is updated via Hebbian plasticity with a learning rate of $\frac{1}{m}$. Upon presentation of token $t$, the matrix $H$ is:

\[H_t = H_{t-1} +  \frac{1}{m}\bell_t \bh_t^T =  H_{t-1} + \frac{1}{m}\bv_t\phi^T(\bk_t) \]
After all the embeddings are passed through the network, assuming $H$ is initially the zero matrix, the final value of $H$ is:

\[H= \frac{1}{m}V\phi^T(K) \]
At the same time, the neuron-to-astrocyte weights are updated via pre-synaptic plasticity. Upon presentation of token $t$, these weights are: 

\[g_{t, i \alpha} = g_{t-1, i \alpha}+ \phi(\biow\bx_t)_\alpha \rightarrow g_{i \alpha} = \sum^{N}_{t = 1}\phi(\bk_t)_{\alpha}\]
Note that as a consequence of the pre-synaptic plasticity, the weight $g_{i \alpha}$ does not depend on the index $i$. Therefore, we will only refer to the vector $\bg \in \fR^{m}$, which--through the pre-synaptic plasticity--is simply the sum over all token presentations of the hidden layer neural activations:

\[\bg = \sum^{N}_{t = 1}\phi(\bk_t)\]

\subsection{Reading Phase}
During the reading phase, the read gate is set to $r = 1$ in \eqref{eq:foward_eqs}, and the inputs are forwarded through the network. The astrocyte process activation value $p$, which according to \eqref{eq:astrocyte_process} does not depend on indices $i$ and $\alpha$, is given by:
\begin{align}
\begin{split}
p = \frac{1}{m}\bh^T\bg = \frac{1}{m}\phi(\bq_t)^T\sum^{N}_{t = 1}\phi(\bk_t) 
\end{split}    
\end{align}
To obtain the last equality we have used $\bh_t = \phi(\biow\bx_t) = \phi(\bq_t)$. Plugging in all the steps of \eqref{eq:foward_eqs}, we see that the last layer has the following value:

\begin{equation}\label{eq:token_wise_approx}
\bell_t = \frac{1}{p}H\phi(\bq_t) + \bx_t =  \frac{V\phi^T(K)\phi(\bq_t)}{\phi(\bq_t)^T\sum^{N}_{t = 1}\phi(\bk_t)} + \bx_t     
\end{equation}
If we compute $\bell$ for every input $\bx_t$ and stack the results column-wise into a matrix $L$, we have:
\begin{equation}\label{eq:bio_out}
L = V\phi^T(K)\phi(Q)G^{-1} + X
\end{equation}
where $G$ is a diagonal matrix such that $G_{zz} = \phi(\bq_z)^T\sum^{N}_{t = 1}\phi(\bk_t)$. 

\begin{figure}[ht]
\begin{center}
\includegraphics[width=1.0\linewidth]{Figures/error_plot.png}
\caption{Error vs number of hidden units ($m$) in our network. As $m$ increases, the difference between the output of the biological circuit and the `pure' Transformer block decreases.}
\label{fig:selfattnvsbio}
\end{center}
\end{figure}

At this point, \eqref{eq:bio_out} is equivalent to a non-autoregressive version of the Linear Transformer \citep{Katharopoulos_Vyas_Pappas_Fleuret_2020}. However, to make our circuit approximate a traditional Transformer with softmax attention, one more piece is required. We need to choose $\phi$ carefully so that we can reproduce the exponential dot-product in the softmax operation. In other words, we need $\phi$ to satisfy: 

\[ \phi(\bx)^T\phi(\mathbf{y}) \approx \text{exp}(\bx^T\mathbf{y}) \]
This can be achieved using random feature attention (RFA) \citep{peng2021random}. RFA relies on a well-known result in kernel approximation theory, which is that the radial basis function (RBF) kernel can, with high probability, be approximated very well using random projections and cosines \citep{rahimi2007random}. To map the RBF kernel to the exponential dot-product kernel used in Transformers, one can simply scale the RBF approximation as follows:

\begin{equation} \label{eq:phi_rbf_approx}
\phi(\bx) = \text{exp}(\frac{||\bx||^2}{2})\phi_{\text{RBF}}(\bx) = \text{exp}(\frac{||\bx||^2}{2})\text{cos}(\Pi\bx + \bb)  
\end{equation}
where the elements of $\Pi \in \fR^{m \times D}$ are drawn from a standard normal distribution, and the elements of $\bb \in \fR^{m}$ are drawn from the uniform distribution on $[0,2\pi]$. The cosine is applied element-wise. Note that we have dropped the usual factor of $\sqrt{\frac{2}{m}}$ from the RBF approximation of  \cite{rahimi2007random}--this is because all constant prefactors of $\phi$ get canceled out in the fraction appearing in \eqref{eq:token_wise_approx}. Plugging \eqref{eq:phi_rbf_approx} into \eqref{eq:bio_out} yields the following:


\begin{align}\label{eq:bio_out_approx}
\begin{split}
L \approx \text{SelfAttn}(X) + X
\end{split}
\end{align}


\subsection{The Role of Normalization and Feedforward Layers}\label{section: normalization}
Thus far we have neglected the role of layer normalization and feedfoward layers. That is because these are operations that are performed on a \textit{per-token} basis, and can therefore be dropped into our model without changing anything in our arguments. In the Transformer model, layer normalization occurs directly preceding and succeeding the FFN layer. With layer normalization and FFN layer, our circuit becomes:

\begin{align*}
\begin{split}
Y = \text{LayerNorm}(L) \approx \text{LayerNorm}(\text{SelfAttn}(X) + X)  \\
\text{Transformer}(X) = \text{LayerNorm}(\text{FFN}(Y) + Y)
\end{split}
\end{align*}
Note that both the FFN and layer norm operations are biologically plausible. Layer normalization is a form of divisive normalization, which is well-studied in the neuroscience literature \citep{carandini2012normalization,Shen_Wang_Navlakha_2021}. In the context of Transformers, it can be achieved by inserting a hidden layer (with lateral connections to implement the normalization) directly before and after the feedforward layer.

\section*{General Case of Untied Weights}\label{section: untied_weights}
In this section we relax the weight tying condition, and generalize our construction to the case when $D \neq d$. Whereas in the previous sections $r$ acted as a gatekeeper for weight matrix $H$, we will now \textit{also} have $r$ act a gatekeeper for a few other weight matrices. Using the same variable names, consider the following neuron-astrocyte forward equations:

\begin{align}\label{eq:foward_eqs_general}
\begin{split}
\beff = \bx & \hspace{.5cm} \in \fR^{d} \\ 
\bh = (1-r)\phi(\biow_K\beff) + r\phi(\biow_Q\beff) & \hspace{.5cm} \in \fR^{m} \\
\bell = r(H\odot\tilde{P})\bh +  (1-r)\biow_V\beff + r\beff    & \hspace{.5cm} \in \fR^{d} \\    
\end{split}
\end{align}
When $r = 0$, we recover the writing phase of \eqref{eq:foward_eqs_write}; when $r=1$, we recover  the reading phase equations of \eqref{eq:token_wise_approx}. When we impose the weight tying constraint of $W_K = W_Q = W$ and $W_V = I$, we recover the original equations of \eqref{eq:foward_eqs}.

\section*{Numerical Validation} The results derived above have also been checked numerically. In Figure \ref{fig:selfattnvsbio} one can see the error between the proposed neuron-astrocyte network and the Transformer block as a function of the ratio of the width of the hidden layer to the size of the token embedding. As expected from the theoretical analysis, the error between the two networks rapidly decreases as the hidden layer becomes wider. In practice, as the width of the hidden layer becomes 5-10 times the embedding dimension the two networks produce very similar outputs.  

\section*{Do We Need Astrocytes?}
Although we are interested in relating Transformers to glial cells, a natural question when positing any new brain mechanism is: ``Can the same behavior be achieved with the old, well-known mechanisms?''. This section demonstrates that an equivalent Transformer circuit can be constructed using just neurons, together with divisive normalization achieved via shunting inhibition. The circuit is similar to \eqref{eq:foward_eqs}:

\begin{align}\label{eq:inhibitory_circuit}
\begin{split}
\beff = \bx & \hspace{.5cm} \in \fR^{d} \\ 
\bh = \phi(\biow\beff) & \hspace{.5cm} \in \fR^{m} \\
R = \bg^T \bh & \hspace{.5cm} \in \fR \\
\bell = \frac{r}{R} H\bh + \beff   & \hspace{.5cm} \in \fR^{d} 
\end{split}
\end{align}
The only difference between \eqref{eq:inhibitory_circuit} and \eqref{eq:foward_eqs} is the addition of a new element, $R$, and the removal of the astrocyte processes. Here $R$ is an inhibitory neuron that divisively normalizes feedfoward inputs into layer $\bell$. However, it does not inhibit all feedfoward inputs equally. Despite both $\bh$ and $\beff$ being feedfoward inputs to layer $\bell$, the divisive inhibition is only implemented on the inputs coming from layer $\bh$. This can happen, for example, if the feedfoward synaptic inputs coming from layer $\bh$ arrive at the dendritic tree close to where inhibitory inputs from neuron $R$ shunt current flow, while the feedfoward inputs coming from layer $\beff$ synapse far away from the shunting \citep{chance2000divisive}. Leaving the reading and writing phases untouched, circuit \eqref{eq:inhibitory_circuit} implements the same forward pass as \eqref{eq:foward_eqs}.


\section*{Time Scale of Transformers in the Brain}
A notable component explicitly missing from Transformers, as well as from our biological circuit, is \textit{time}. Unlike the brain, Transformers are not dynamical systems. Except in the special instance of autoregressive linear Transformers (which may be seen as a kind of recurrent neural network), Transformers do not maintain a hidden state via recurrent weights. Nevertheless, timing is a critical aspect of brain processing, so in this section we describe a plausible timescale for our circuit in terms of known biophysics parameters. The following paragraph argues that the total compute time for our circuit is on the order of $(N+1) \times 100$ milliseconds, where $N$ is the number of tokens. 

Our circuit operates in two distinct phases: a reading phase and a writing phase. The reading phase does not involve any plasticity, so the only relevant timescale to compute is how long it takes to traverse the neuron $\rightarrow$ astroctyte $\rightarrow$ synapse pathway. Recent data indicate that this entire signalling pathway can be traversed on the order of a few hundred milliseconds \citep{stobart2018cortical}. The speed of the writing phase limited by the speed of plasticity. There are two types of plasticity used in our model: 1) Hebbian plasticity between neurons and 2) pre-synaptic plasticity between neurons and astrocyte processes. The former can happen on relatively fast timescale, for example through spike-timing-dependent plasticity (STDP). In that case of STDP, the timescale can be as fast as a few tens of milliseconds \citep{klampfl2013emergence}.  The latter timescale is harder to establish, due to limitations in calcium recording technology. While fast calcium transients in astrocyte processes have been recently recorded \citep{stobart2018cortical}, and neuron-astrocyte plasticity has been experimentally observed \citep{croft2015plasticity}, \textit{fast} (e.g < 1s) neuron-astrocyte plasticity has not been observed. Assuming the timescale of this plasticity is also a few hundred milliseconds, then an  estimate for the overall timescale of our model would be a few hundred milliseconds for the read phase, plus a few hundred milliseconds \textit{per token} for the write phase. This leads to a total compute time of $\tau_{compute} = (100 + N \times 100)$ milliseconds. Finally, note that most "real-world" Transformers are composed of many blocks. Our circuit is a model of a single block. Thus, the total compute time for a $k$ block transformer modelled using our framework is $k\tau_{compute}$.

\section*{Discussion}
Our paper serves a dual purpose. The first is to provide a simple, biological account of how a Transformer might be implemented using brain-like elements. The second is to propose a specific, functional role for glial cells as normalization units in the self-attention operator. While the feedback loop between neurons and glia is well-studied from a biological perspective, there is comparatively little work studying it from a computational perspective \citep{kastanenka2020roadmap}. Most of the computational studies (with notable exceptions, see e.g \citep{song2016design,tang2019introducing,ivanov2021increasing,Peterson_2022}) are focused on understanding the biophysics of neuron-glia or glia-glia signalling, as opposed to the abstracted functions of neuron-glia circuits \citep{de2009glutamate,de2011tale,goldberg2010nonlinear,de2012computational}. Our paper proposes a potential role for glial cells along the lines presented in \cite{kastanenka2020roadmap}. In particular, we propose that astrocytes are involved in attention via their role in normalization literally.


Finally, as calcium imaging technology improves, it will become increasingly feasible to compare artificial representations in neural networks to biological representations in astrocyte networks (as has been done for biological neural networks). Our work suggests that a reasonable place to start may be to compare Transformer normalization terms to astrocytic calcium responses.

\begin{acknowledgements}
This work was completed during the MIT-IBM Watson AI Lab Summer 2022 Internship Program. We thank John Hopfield and Mriganka Sur for helpful comments and feedback. 
\end{acknowledgements}

\section*{Bibliography}
\bibliography{bio_transformer}

\end{document}
