## Supplementary material for "Building Transformers from Neurons and Astrocytes": LaTeX files: preamble.tex

%fancy R
\newcommand{\fR}{\mathbb{R}}

%number of elements in sequence
\newcommand{\slen}{N}

%bold x
\newcommand{\bx}{\mathbf{x}}

%bold v
\newcommand{\bv}{\mathbf{v}}

%bold k
\newcommand{\bk}{\mathbf{k}}

%bold q
\newcommand{\bq}{\mathbf{q}}

%bold g
\newcommand{\bg}{\mathbf{g}}

%
\newcommand{\bell}{\boldsymbol{\ell}}

%
\newcommand{\beff}{\mathbf{f}}

%
\newcommand{\bh}{\mathbf{h}}

%
\newcommand{\ba}{\mathbf{a}}

%
\newcommand{\bb}{\mathbf{b}}

%dimension of embedding vector
\newcommand{\edim}{d}

%bio_circuit weight
\newcommand{\biow}{W}

%calcium
\newcommand{\ca}{\text{Ca}^{2+}}

%softmax function
\newcommand{\softmax}{\text{softmax}}

%big derivative dot
\newcommand*{\dt}[1]{\accentset{\mbox{\large\bfseries .}}{#1}}

%trace
\DeclareMathOperator{\Tr}{Tr}
