## Supplementary figures and images for "Building Transformers from Neurons and Astrocytes"

### error_plot.png

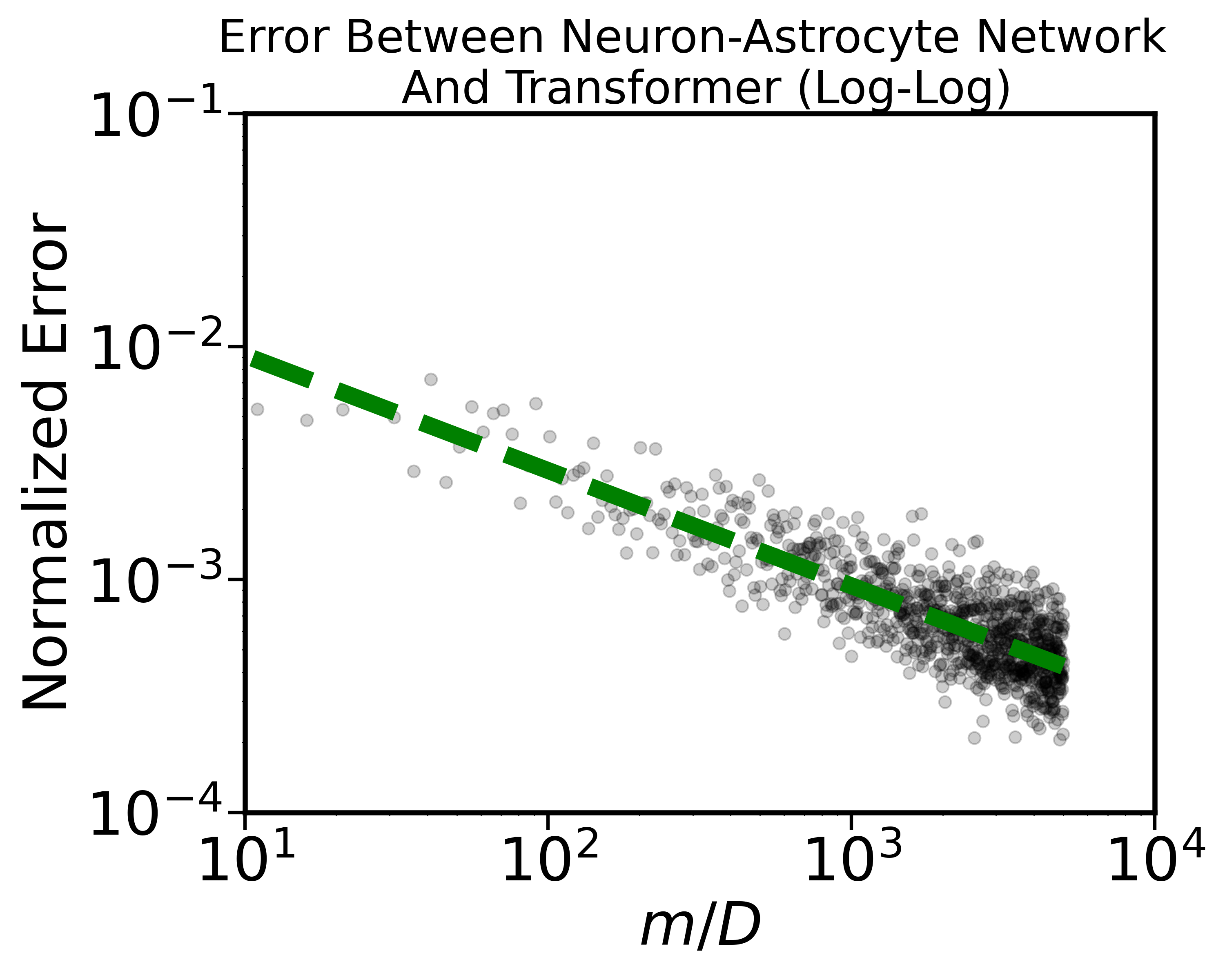

### pnas_fig_1_long_version_2.png

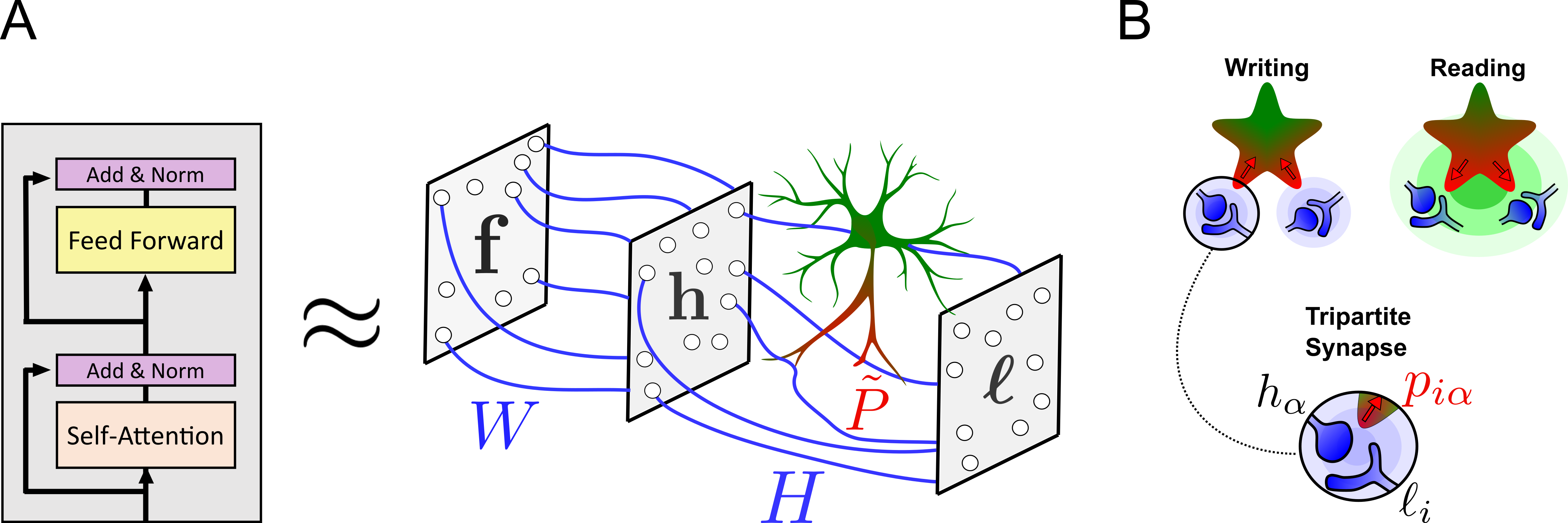
